# Resolving the Dynamic Motions of SARS-CoV-2 nsp7 and nsp8 Proteins Using Structural Proteomics

**DOI:** 10.1101/2021.03.06.434214

**Authors:** Valentine V. Courouble, Sanjay Kumar Dey, Ruchi Yadav, Jennifer Timm, Jerry Joe E. K. Harrison, Francesc X. Ruiz, Eddy Arnold, Patrick R. Griffin

## Abstract

Coronavirus (CoV) non-structural proteins (nsps) assemble to form the replication-transcription complex (RTC) responsible for viral RNA synthesis. nsp7 and nsp8 are important cofactors of the RTC, as they interact and regulate the activity of RNA-dependent RNA polymerase (RdRp) and other nsps. To date, no structure of full-length SARS-CoV-2 nsp7:nsp8 complex has been published. Current understanding of this complex is based on structures from truncated constructs or with missing electron densities and complexes from related CoV species with which SARS-CoV-2 nsp7 and nsp8 share upwards of 90% sequence identity. Despite available structures being solved using crystallography and cryo-EM representing detailed snapshots of the nsp7:nsp8 complex, it is evident that the complex has a high degree of structural plasticity. However, relatively little is known about the conformational dynamics of the complex and how it assembles to interact with other nsps. Here, the solution-based structural proteomic techniques, hydrogen-deuterium exchange mass spectrometry (HDX-MS) and crosslinking mass spectrometry (XL-MS), illuminate the structural dynamics of the SARS-CoV-2 full-length nsp7:nsp8 complex. The results presented from the two techniques are complementary and validate the interaction surfaces identified from the published three-dimensional heterotetrameric crystal structure of SARS-CoV-2 truncated nsp7:nsp8 complex. Furthermore, mapping of XL-MS data onto higher order complexes suggests that SARS-CoV-2 nsp7 and nsp8 do not assemble into a hexadecameric structure as implied by the SARS-CoV full-length nsp7:nsp8 crystal structure. Instead our results suggest that the nsp7:nsp8 heterotetramer can dissociate into a stable dimeric unit that might bind to nsp12 in the RTC without altering nsp7-nsp8 interactions.

## INTRODUCTION

The severe acute respiratory syndrome coronavirus 2 (SARS-CoV-2; CoV-2) is the agent responsible for the coronavirus disease 2019 (COVID-19) that has infected over 110 million people and claimed over 2.4 million lives worldwide [1, 2]. As part of the betacoronavirus genus, CoV-2 is an enveloped positive single-stranded RNA virus [3]. The large ∼30 kb genome encodes six functional open reading frames (ORF): replicase (ORF1a/ORF1b), spike (S), envelope (E), membrane (M), and nucleocapsid (N), ordered from the 5’ to 3’ end. An additional seven putative ORFs encoding accessory proteins are interspersed between the structural proteins [4].

Replication of the very large genome is mediated by a highly dynamic protein-RNA complex known as the replication-transcription complex (RTC) [5]. The non-structural proteins (nsps) that make up this complex are encoded by the replicase genes. ORF1a and ORF1b, which take up approximately two thirds of the viral genome, are translated to produce two large polyproteins, pp1a and pp1ab, which then undergo proteolytic processing to release 16 individual nsps. This process is mediated by two self-encoded proteases: the papain-like protease (PLpro; nsp3) responsible for cleaving nsp1-4 and the main chymotrypsin-like protease (Mpro; 3CLpro; nsp5) responsible for cleaving nsp4-16.

Over the past year, many studies have focused on solving the structure of the RNA-polymerase complex to aid in the development of antiviral drugs to treat CoV-2 infections [6-11]. These structures suggest that the minimal RNA-polymerase complex is composed of the RNA-dependent RNA-polymerase (RdRp) catalytic domain encoded in the C-terminus of nsp12 bound to two cofactor proteins nsp7 and nsp8. Cryo-electron microscopy (cryo-EM) structures have suggested the stoichiometry of nsp12:nsp7:nsp8 for the complex to be 1:1:2, respectively. *In vitro* studies have shown that the polymerase activity of nsp12 requires binding of at least nsp7 and nsp8. Structural and biochemical analysis of the nsp7 and nsp8 complex in the absence of nsp12 have been less studied.

There are currently seven available crystal structures for the CoV-2 nsp7-nsp8 complex in the Protein Data Bank (PDB) [12-18]. Only one of these structures has been peer-reviewed and published [12]. Overall, the seven structures can be classified into three representative conformations: A) dimer (PDB: 6WIQ and 6M5I), B) linear heterotetramer (PDB: 6YHU, 6WQD, and 7JLT) and C) cubic heterotetramer (PDB: 6WTC and 6XIP). Comparison of the heterotetrameric structures reveals that the linear heterotetramer structures are arranged around a nsp7 dimer core while the cubic heterotetramer structures are composed of two adjacent nsp7:nsp8 dimers. Interestingly, X-ray crystal structures of full-length nsp7:nsp8 complex from SARS-CoV and from feline coronavirus reveal alternative conformations. The SARS-CoV complex forms a hexadecameric structure primarily organized around N-terminal nsp8 interactions [19]. On the contrary, the feline coronavirus structure revealed a nsp7:nsp8 heterotrimer with one nsp8 molecule associated with two nsp7 molecules that self-interact [20]. Furthermore, a study of SARS-CoV nsp7:nsp8 complex using native mass spectrometry (MS) reported a heterotetrameric structure with no nsp7 self-interaction [21]. Nevertheless, to date the structure of the intact full length nsp7:nsp8 complex from CoV-2 has not been reported. To address this lack of knowledge, we applied a combination of hydrogen-deuterium exchange MS (HDX-MS) and crosslinking MS (XL-MS) to probe the dynamics of the intact full length nsp7:nsp8 complex from CoV-2. Our studies demonstrate that the nsp7:nsp8 complex contains a high degree of architectural plasticity that is not apparent from X-ray crystallographic or cryo-EM studies. Results from the structural proteomic studies are used to propose a new model of assembly.

## EXPERIMENTAL

### Reagents and Plasmids

Unless otherwise specified, all chemical and reagents were purchased from Sigma-Aldrich (St. Louis, MO). Formic acid, trifluoroacetic acid, and UHPLC-grade solvents were purchased from ThermoFisher. pGBW-m4046979 (coding for full-length nsp7, NCBI Reference Sequence: YP_009725303.1, codon optimized, with an initial Met and a cleavable C-terminal TEV His_6_ tag) was a gift from Ginkgo Bioworks (Addgene plasmid # 145611; http://n2t.net/addgene:145611; RRID:Addgene_145611). pGBW-m4046852 (coding for full-length nsp8, NCBI Reference Sequence: YP_009725304.1, codon optimized, with an initial Met and a cleavable C-terminal TEV His_6_ tag) was a gift from Ginkgo Bioworks (Addgene plasmid # 145584; http://n2t.net/addgene:145584; RRID:Addgene_145584).

### Protein Expression and Purification

nsp7 was expressed in T7 express competent cells (previously transformed with the aforementioned plasmid) at 37°C, then when OD_600_ ∼0.6 the culture was brought to 20°C for 1 h, followed by overnight induction with 1 mM IPTG at 20°C. Cells were pelleted by centrifugation at 7,200 x g and after a freeze-thaw cycle, were resuspended with lysis buffer containing 50 mM Tris pH 8.0, 500 mM NaCl, 20 mM imidazole, 10 mM CHAPS, 5% glycerol, 1 mM TCEP, 1 µM leupeptin, 1 µM pepstatin A, 1 mM PMSF, sonicated and centrifugated at 30,000 x g for 60 min. Ni-NTA affinity purification was performed, and after extensive wash with lysis buffer, an imidazole-based step elution was done, followed by overnight digestion with TEV protease and a reverse Ni-NTA affinity purification to recover the cleaved protein. Subsequent heparin affinity and Mono Q anion exchange steps further improved the purity of nsp7 protein. nsp8 was similarly expressed, with an additional size exclusion chromatography step was done after the heparin affinity and Mono Q steps. The final buffer for both proteins was 50 mM HEPES pH 8.0, 500 mM NaCl, 5% glycerol, 1 mM TCEP.

### Crosslinking mass spectrometry (XL-MS)

#### Sample preparation

For DSSO (disuccinimidyl sulfoxide) (ThermoFisher) crosslinking reactions, individual protein and protein-protein complexes were diluted to 10 µM in crosslinking buffer (50 mM HEPES pH 8.0, 500 mM NaCl, 1 mM TCEP) and incubated for 30 min at room temperature prior to initiating the crosslinking reaction. DSSO crosslinker was freshly dissolved in crosslinking buffer to a final concentration of 75 mM before being added to the protein solution at a final concentration of 1.5 mM. The reaction was incubated at 25°C for 45 min, then quenched by adding 1 µL of 1.0 M Tris pH 8.0 and incubating an additional 10 min at 25°C. Control reactions were performed in parallel without adding the DSSO crosslinker. All crosslinking reactions were carried out in three replicates. The presence of crosslinked proteins was confirmed by comparing to the no crosslink negative control samples using SDS-PAGE and Coomassie staining. The remaining crosslinked and non-crosslinked samples were separately pooled and then precipitated using methanol and chloroform. Dried protein pellets were resuspended in 12.5 µL of resuspension buffer (50 mM ammonium bicarbonate, 8M urea, pH 8.0). ProteaseMAX (Promega - V5111) was added to 0.02% and the solutions were mixed on an orbital shaker operating at 400 RPM for 5 min. After resuspension, 87.5 µL of digestion buffer (50 mM ammonium bicarbonate, pH 8.0) was added. Protein samples were reduced by adding 1 µL of 500 mM DTT followed by incubation of the protein solutions in an orbital shaker operating at 400 RPM at 56°C for 20 minutes. After reduction, 2.7 µL of 550 mM iodoacetamide was added and the solutions were incubated at room temperature in the dark for 15 min. Reduced and alkylated protein solutions were digested overnight using trypsin at a ratio of 1:150 (w/w trypsin:protein) at 37°C. Peptides were acidified to 1% trifluoroacetic acid (TFA) and then desalted using C_18_ ZipTip® (Millipore cat # ZTC18 5096). Dried peptides were resuspended in 10 µL of 0.1% TFA in water. Samples were then frozen and stored at −20°C until LC-MS analysis.

#### Liquid Chromatography and Mass Spectrometry

500 ng of sample was injected (triplicate injections for crosslinked samples and duplicate injections for control samples) onto an UltiMate 3000 UHP liquid chromatography system (Dionex, ThermoFisher). Peptides were trapped using a μPAC C18 trapping column (PharmaFluidics) using a load pump operating at 20 µL/min. Peptides were separated on a 200 cm μPAC C18 column (PharmaFluidics) using a linear gradient (1% Solvent B for 4 min, 1-30% Solvent B from 4-70 min, 30-55% Solvent B from 70-90 min, 55-97% Solvent B from 90-112 min, and isocratic at 97% Solvent B from 112-120 min) at a flow rate of 800 nL/min. Gradient Solvent A contained 0.1% formic acid and Solvent B contained 80% acetonitrile and 0.1% formic acid. Liquid chromatography eluate was interfaced to an Orbitrap Fusion Lumos Tribrid mass spectrometer (ThermoFisher) with a Nanospray Flex ion source (ThermoFisher). The source voltage was set to 2.5 kV, and the S-Lens RF level was set to 30%. Crosslinks were identified using a previously described MS2-MS3 method [22] with slight modifications. Full scans were recorded from *m*/*z* 150 to 1500 at a resolution of 60,000 in the Orbitrap mass analyzer. The AGC target value was set to 4e5, and the maximum injection time was set to 50 ms in the Orbitrap. MS2 scans were recorded at a resolution of 30,000 in the Orbitrap mass analyzer. Only precursors with charge state between 4-8 were selected for MS2 scans. The AGC target was set to 5e4, a maximum injection time of 150 ms, and an isolation width of 1.6 m/z. CID fragmentation energy was set to 25%. The two most abundant reporter doublets from the MS2 scans with a charge state of 2-6, a 31.9721 Da mass difference (Kao et al 2011), and a mass tolerance of ±10 ppm were selected for MS3. The MS3 scans were recorded in the ion trap in rapid mode using HCD fragmentation with 35% collision energy. The AGC target was set to 2e4, the maximum injection time was set for 200 ms, and the isolation width set to 2.0 m/z.

#### Data Analysis

To identify crosslinked peptides, Thermo.Raw files were imported into Proteome Discoverer 2.5 (ThermoFisher) and analyzed via XlinkX algorithm [23] using the MS2_MS3 workflow with the following parameters: MS1 mass tolerance—10 ppm; MS2 mass tolerance—20 ppm; MS3 mass tolerance—0.5 Da; digestion—trypsin with four missed cleavages allowed; minimum peptide length of 4 amino acids, fixed modification—carbamidomethylation (C); variable modification—oxidation (M); and DSSO (K, S, T, Y). The XlinkX/PD Validator node was used for crosslinked peptide validation with a 1% false discovery rate (FDR). Identified crosslinks were further validated and quantified using Skyline (version 19.1) [24] using a previously described protocol [25]. Crosslink spectral matches found in Proteome Discoverer were exported and converted to sequence spectrum list format using Excel (Microsoft). Crosslink peak areas were assessed using the MS1 full-scan filtering protocol for peaks within 8 min of the crosslink spectral match identification. Peak areas were assigned to the specified crosslinked peptide identification if the mass error was within 10 ppm of the theoretical mass, the isotope dot product was greater than 0.95, and if the peak was not found in the non-crosslinked negative control samples. The isotope dot product compares the distribution of the measured MS1 signals against the theoretical isotope abundance distribution calculated based on the peptide sequence. Its value ranges between 0 and 1, where 1 indicates a perfect match [26]. Pair-wise comparisons were made using ‘MSstats’ package [27] implemented in Skyline to calculate relative fold changes and significance. Significant change thresholds were defined as a log_2_ fold change less than −2 or greater than 2 and -log_10_ p-value greater than 1.3 (p-value less than 0.05). Visualization of proteins and crosslinks was generated using xiNET [28].

### Hydrogen-deuterium exchange mass spectrometry (HDX-MS)

Solution-phase amide HDX experiments were carried out with a fully automated system (CTC HTS PAL, LEAP Technologies, Carrboro, NC; housed inside a 4°C cabinet) as previously described [29] with the following modifications. Peptides were identified using tandem MS (MS/MS) experiments performed on a QExactive (Thermo Fisher Scientific, San Jose, CA) over a 70 min gradient. Product ion spectra were acquired in a data-dependent mode and the five most abundant ions were selected for the product ion analysis per scan event. The MS/MS *.raw data files were converted to *.mgf files and then submitted to MASCOT (version 2.3 Matrix Science, London, UK) for peptide identification.

The maximum number of missed cleavages was set at 4 with the mass tolerance for precursor ions +/-0.6 Da and for fragment ions +/-8 ppm. Oxidation to methionine was selected for variable modification. Pepsin was used for digestion and no specific enzyme was selected in MASCOT during the search. Peptides included in the peptide set used for HDX detection had a MASCOT score of 20 or greater. The MS/MS MASCOT search was also performed against a decoy (reverse) sequence and false positives were ruled out if they did not pass a 1% false discovery rate.

For differential HDX, protein-protein complexes were formed by incubating nsp7 and nsp8 at 1:1, 3:1, or 1:3 molar ratios for 30 min at room temperature. The reactions (5 μL) were mixed with 20 μL of D_2_O-containing HDX buffer (50 mM HEPES, 500 mM NaCl, 1 mM TCEP, pD 8.4) and incubated at 4°C for 0 s, 10 s, 30 s, 60 s, 900 s or 3600 s. Following on-exchange, unwanted forward-or back-exchange was minimized, and the protein was denatured by the addition of 25 μL of a quench solution (5 M Urea, 1% TFA, pH 2). Samples were then immediately passed through an immobilized pepsin column (prepared in house) at 50 μL min^-1^ (0.1% v/v TFA, 4°C) and the resulting peptides were trapped and desalted on a 2 mm × 10 mm C_8_ trap column (Hypersil Gold, ThermoFisher). The bound peptides were then gradient-eluted (4-40% CH_3_CN v/v and 0.3% v/v formic acid) across a 2.1 mm × 50 mm C_18_ separation column (Hypersil Gold, ThermoFisher) for 5 min. Sample handling and peptide separation were conducted at 4°C. The eluted peptides were then subjected to electrospray ionization directly coupled to a high resolution Orbitrap mass spectrometer (QExactive, ThermoFisher). Each HDX experiment was injected in triplicate. The intensity weighted mean m/z centroid value of each peptide envelope was calculated and subsequently converted into a percentage of deuterium incorporation. This is accomplished by determining the observed averages of the undeuterated and fully deuterated spectra using the conventional formula described elsewhere [30]. The fully deuterated control, 100% deuterium incorporation, was calculated theoretically, and corrections for back-exchange were made on the basis of an estimated 70% deuterium recovery and accounting for 80% final deuterium concentration in the sample (1:5 dilution in D_2_O HDX buffer). Statistical significance for the differential HDX data is determined by an unpaired t-test for each time point, a procedure that is integrated into the HDX Workbench software [31].

The HDX data from all overlapping peptides were consolidated to individual amino acid values using a residue averaging approach. Briefly, for each residue, the deuterium incorporation values and peptide lengths from all overlapping peptides were assembled. A weighting function was applied in which shorter peptides were weighted more heavily and longer peptides were weighted less. Each of the weighted deuterium incorporation values were then averaged incorporating this weighting function to produce a single value for each amino acid. The initial two residues of each peptide, as well as prolines, were omitted from the calculations. This approach is similar to that previously described [32].

HDX analyses were performed in triplicate. Deuterium uptake for each peptide is calculated as the average of %D for all on-exchange time points and the difference in average %D values between the unbound and bound samples is presented as a heat map with a color code given at the bottom of the figure (warm colors for deprotection and cool colors for protection). Peptides are colored by the software automatically to display significant differences, determined either by a >5% difference (less or more protection) in average deuterium uptake between the two states, or by using the results of unpaired t-tests at each time point (p-value < 0.05 for any two time points or a p-value < 0.01 for any single time point). Peptides with non-significant changes between the two states are colored grey. The exchange at the first two residues for any given peptide is not colored. Each peptide bar in the heat map view displays the average Δ %D values, associated standard deviation, and the charge state. Additionally, overlapping peptides with a similar protection trend covering the same region are used to rule out data ambiguity.

## RESULTS AND DISCUSSION

### HDX-MS and XL-MS are complementary techniques to investigate structural dynamics

While X-ray crystallography and cryo-EM are the gold-standard techniques for determining atomic resolution structures of macromolecular complexes, they represent snapshots of protein structures that may reveal some aspects of conformational ensembles, but do not fully reflect the structural plasticity inherent to proteins in solution. Characterizing this structural plasticity is essential to understand the structure and function relationship of proteins and protein complexes. Hydrogen-deuterium exchange mass spectrometry (HDX-MS) and crosslinking MS (XL-MS) are complementary techniques that can be used to garner information on the solution phase dynamics of proteins and protein complexes. Specifically, HDX-MS reports on protein backbone dynamics while XL-MS reports on side-chain residency and reactivity.

### HDX-MS provides a footprint of nsp7 and nsp8 secondary structure

HDX-MS measures changes in mass due to isotopic exchange of the amide hydrogens of the protein backbone with the solvent. The rate of isotopic exchange is influenced by the intrinsic properties of the amino acid sequence, the folded state of the protein, and the dynamics of the hydrogen-bonding network. Given that protein secondary structures such as α-helices and β-sheets are characterized and stabilized via hydrogen bonds, HDX-MS can be used to predict or assess the dynamics of proteins’ secondary structure elements. As shown in **Figure 1A and 1B**, HDX-MS analysis of nsp7 and nsp8 proteins in isolation reveals solvent exchange behavior that is largely in agreement with the secondary structure observed in the published nsp7:nsp8 heterotetrameric crystal structure (PDB:6YHU) [12]. For nsp7, the three α-helical bundles within the protein (H1^nsp7^ Lys2-Leu20, H2^nsp7^ Ser26-Leu41, H3^nsp7^ Thr45-Ser61) are characterized by lower deuterium uptake relative to other regions of the protein. For nsp8, the C-terminal subdomain is predicted to be composed of four antiparallel β-strands (B1^nsp8^ Ala125-Ile132, B2^nsp8^ Thr146-Tyr151, B3^nsp8^ Ala152-Val160, B4^nsp8^ Leu186-Arg190) with an α-helix inserted between the first two β-strands (H3^nsp8^ Tyr135-Thr142). The α-helical region of this domain has the lowest deuterium uptake. However, the other two predicted α-helices (H1^nsp8^ Glu77-Leu98, H2^nsp8^ Asp101-Asp112) show high percent deuterium uptake suggesting greater conformational dynamics such that these regions may not actually be ordered into helices when nsp8 is in isolation.

**Figure 1.**
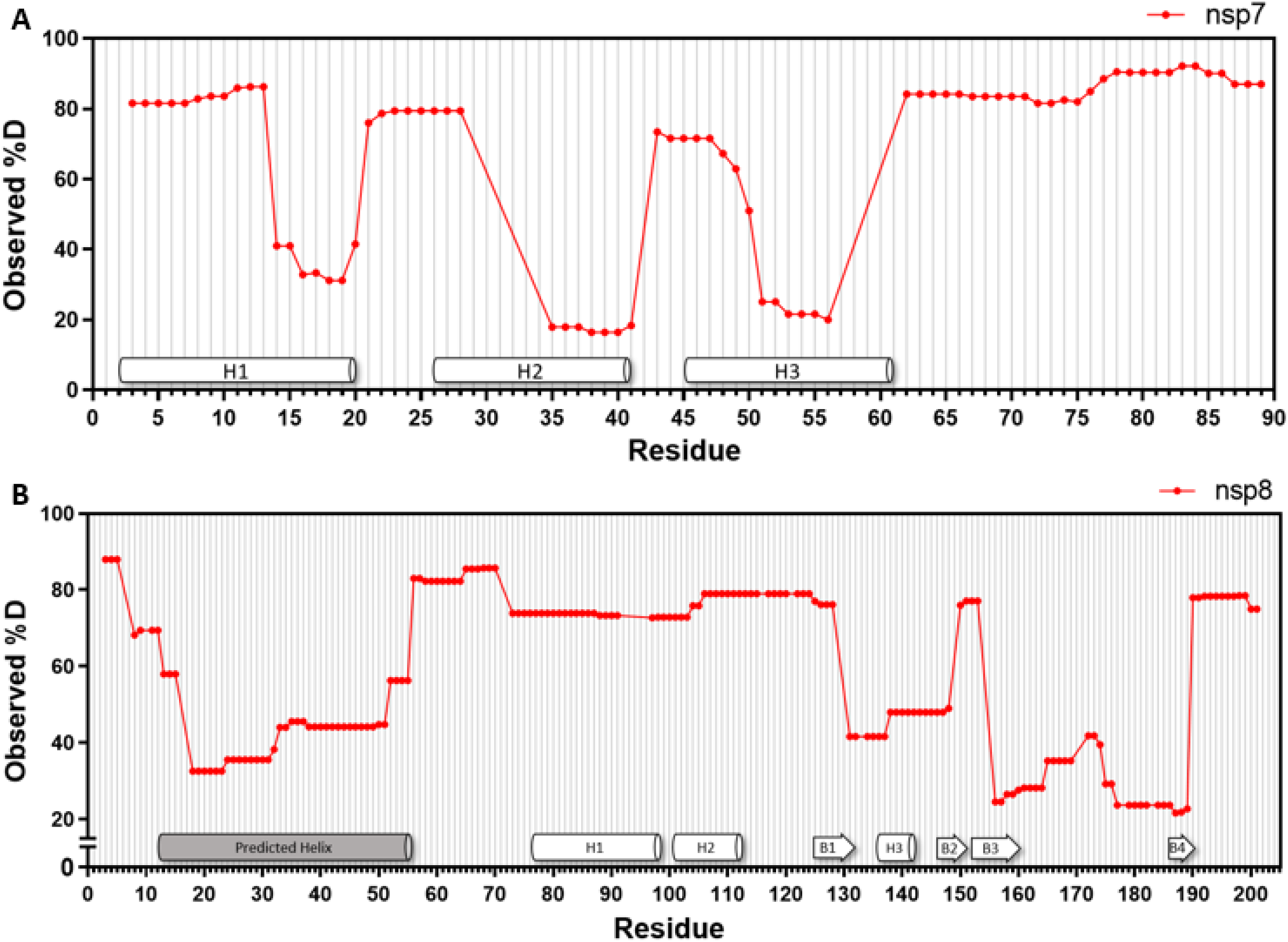
HDX-MS analysis of nsp7, nsp8, and nsp7:nsp8 complexes. Observed percent deuterium uptake of nsp7 **(A)** and nsp8 **(B)** during the 1h experiment time course. Secondary structure from PDB: 6YHU annotated in white and predicted secondary structure based on observed percent deuterium in gray.

As the X-ray structure (PDB: 6YHU) used truncated forms of both nsp7 and nsp8 to facilitate crystallization, the HDX footprint of the full-length protein can predict secondary structure of the missing regions. The truncated nsp8 N-terminus (residues 1-75, note that residues 193-198 in the C-terminus are truncated too) shows low deuterium uptake, 30-60% observed deuterium which suggests the presence of helical structure in this region. This prediction is supported by available cryo-EM structures of nsp8 in complex with nsp7, nsp12, and RNA that reveal the nsp8 N-terminus to be primarily helical in nature [7-10]. This region is not always fully resolved in the absence of RNA suggesting that it may be more mobile and sampling multiple orientations, which is supported by the HDX results showing that this region has somewhat higher percent deuterium uptake (32-45% deuterium uptake) compared to the other helices in nsp8 and nsp7 (20-25% deuterium uptake). The truncated C-terminus of nps7 (residues 72-83) shows high levels of deuterium uptake suggesting greater dynamics in this region and potentially lacking defined secondary structure (**Figure 4**).

### Differential HDX validates nsp7:nsp8 binding surfaces

Differential HDX (comparison of the exchange kinetics of individual proteins in isolation to that within the protein complex) of the nsp7:nsp8 complex was initially conducted at a 1:1 molar ratio in line with the reported stoichiometry [12]. Reduced deuterium uptake was only observed in nsp8 in H2^nsp8^, B1^nsp8^, and B3^nsp8^ (**Figure 2A, S1A, S2A, Table S1**). Based on the X-ray structure (PDB: 6YHU), H2^nsp8^ is located at the nsp7-nsp8 dimerization interface. This interface is stabilized by a leucine zipper motif composed of nsp7 Leu56, Leu60, and Leu71 and nsp8 Leu95 and Leu102. Regions within nsp8 containing both Leu residues are significantly protected to exchange when nsp8 is in complex with nsp7. Furthermore, peptides containing nsp8 Phe92 and adjacent hydrophobic residues implicated in stabilizing the complex are significantly protected from exchange [12]. Surprisingly, there was no detectable protection to solvent exchange in nsp7 under the experimental conditions employed at a 1:1 molar ratio (**Figure 2B, S1B, S2B, Table S2**). Therefore, we repeated the analysis with 3-fold molar excess of nsp8 to saturate all nsp7 binding sites (**Figure 2D, S2C, S2B, Table S2**). In this experiment, with excess nsp8, protection to exchange was observed in all three α-helices of nsp7 as well as in its C-terminal region. H1^nsp7^ and H3^nsp7^ are predicted to be involved in forming the nsp7:nsp8 dimer interface while H1^nsp7^ and H2^nsp7^ are predicted to be involved with the nsp7:nsp8 2:2 heterotetrameric interface [12]. Significant protection from solvent exchange in H1^nsp7^ as well as the N-termini of H2^nsp7^ and H3^nsp7^ are consistent with this speculation. Regions of nsp7 that contain a disulfide bridge formed between the symmetric nsp7 Cys8 residues to stabilize the heterotetramer demonstrated the highest magnitude of solvent protection. Interestingly, the C-terminal region of nsp7 showed levels of protection similar to that of the nsp7 Cys8 region. In PDB: 6YHU this region is not observed and thus was not reported to be involved in the interaction with nsp8. However, the other structures in the linear heterotetramer group (PDB:6WQD and PDB:7JLT) were able to resolve more of the C-termini of nsp7 and these structures show that this region is organized into a fourth α-helix (D67-R79). The significant protection to solvent exchange observed in this region supports the formation of H4^nsp7^ upon binding to nsp8 (**Table S2**).

**Figure 2.**
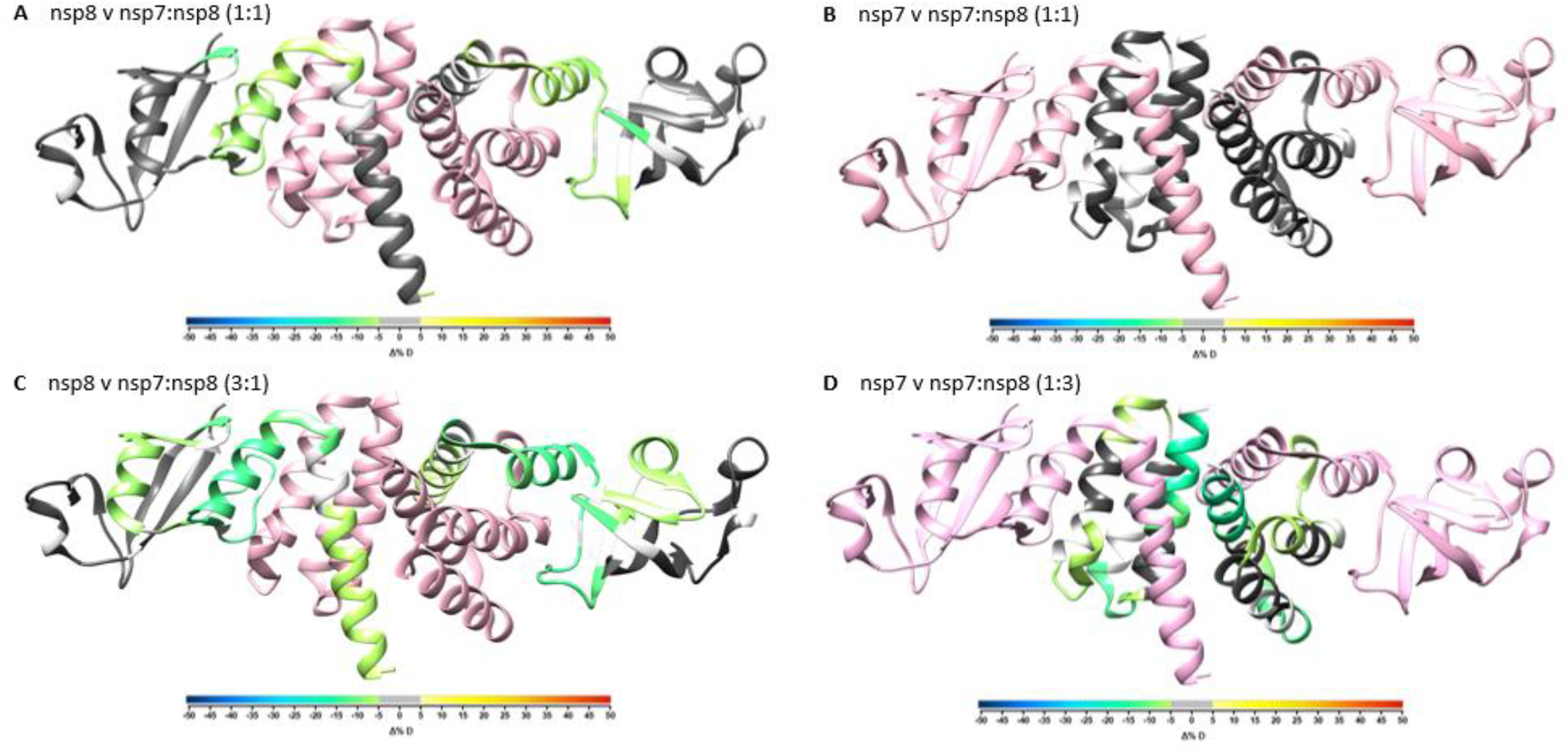
Overlay of differential HDX-MS perturbation values onto PDB:6YHU. Perturbation values for nsp7 vs nsp7:nsp8 1:1 **(A)**, nsp8 vs nsp7:nsp8 1:1 **(B)**, nsp7 vs nsp7:nsp8 1:3 **(C)**, and nsp8 vs nsp7:nsp8 3:1 **(D)** colored according to change in percent deuterium levels shown in color bar and respective partner protein colored in pink. Residues not observed by HDX-MS are colored in white.

To complement the above experiment, we repeated the differential HDX analysis of nsp8 with a 3-fold molar excess of nsp7 to saturate nsp8 binding sites (**Figure 1D, S1D**). In the presence of excess nsp7, increased magnitude of protection to solvent exchange was observed in regions that showed protection from exchange at 1:1 molar ratio. Additional protection to exchange was observed in new regions including H1^nsp8^, H3^nps8^, and B2^nsp8^. The HDX analysis of nsp8 in complex with nsp7 suggest that observed H1^nsp8^ and H2^nps8^ in the nsp7:nsp8 crystal structure are only formed upon binding to nsp7 as this region shows high percent deuterium levels in isolation and significantly lower percent deuterium levels in the presence of nsp7.

### XL-MS validates nsp7:nsp8 direct interaction

To complement the analysis of backbone dynamics, we conducted differential XL-MS to observe changes in side-chain residency and reactivity. Figure S3 shows the XL reaction efficiency assessed by SDS-PAGE. To minimize false assignment of crosslink products we utilized the MS cleavable crosslinker disuccinimidyl sulfoxide (DSSO). Crosslinked peptides are cleaved using collision-induced dissociation (CID) at the MS^2^ level to produce unique fragment pair ions that trigger MS^3^ analysis of the single peptide chain fragments allowing for unambiguous sequence identification. Crosslinks were identified as described in the Experimental section using the XlinkX software in Proteome Discoverer [23] and crosslink spectral matches were exported for further manual validation in Skyline [24] using the MS1 full-scan filtering protocol. Peak areas were only assigned to crosslinked peptides if the isotope dot product was greater than 0.95 and if the peak was not found in the non-crosslinked negative control sample. This allowed us to perform differential XL-MS via relative pair-wise comparison using the MSstats package in Skyline [27]. Threshold for significant changes in crosslinks were set to a log_2_ fold change less than −2 or greater than 2 and -log_10_ p-value greater than 1.3 (p-value less than 0.05).

XL-MS of nsp7 in isolation resulted in no detectable intra-protein crosslinks. Several dead-end crosslinks were identified (**Figure S4A**). The presence of only dead-end crosslinks suggests nsp7 to be in a more linear conformation unable to form intra-protein crosslinks. It also suggests no significant nsp7:nsp7 side-chain interactions (homomultimerization). While there have been mixed reports suggesting that nsp7 does or does not engage in self-interaction [33, 34], the results presented here support that nsp7 does not self-interact. Interestingly, in the presence of nsp8, the residues with dead-end crosslinks all form inter-nsp7-nsp8 crosslinks with nsp8. XL-MS analysis of nsp8 in isolation resulted in the identification of 25 intra-protein crosslinks (**Figure S4B**). As shown in Figure 3A, XL-MS analysis of the nsp7:nsp8 complex resulted in the identification of both intra-nsp8 crosslinks as well as inter-nsp7-nsp8 crosslinks. A total of four inter-nsp7-nsp8 crosslinks were identified in the nsp7:nsp8 complex at a 1:1 molar ratio. Comparing crosslinks detected in nsp8 to those detected in the nsp7:nsp8 complex, we observed that a majority of the intra-nsp8 crosslinks are not significantly different (unchanged) between the two sample groups. However, the intra-nsp8 crosslinks located near Lys79 which forms inter-nsp7-nsp8 crosslinks in the nsp7:nsp8 sample group, are significantly enriched in the nsp8 alone samples (decreased in the nsp7:nsp8 complex). These data suggest that residues adjacent to nsp8 Lys79 are involved in the interaction with nsp7 such that formation of inter-nsp7-nsp8 crosslinks is favored over intra-nsp8 crosslinks when in the presence of nsp7.

**Figure 3.**
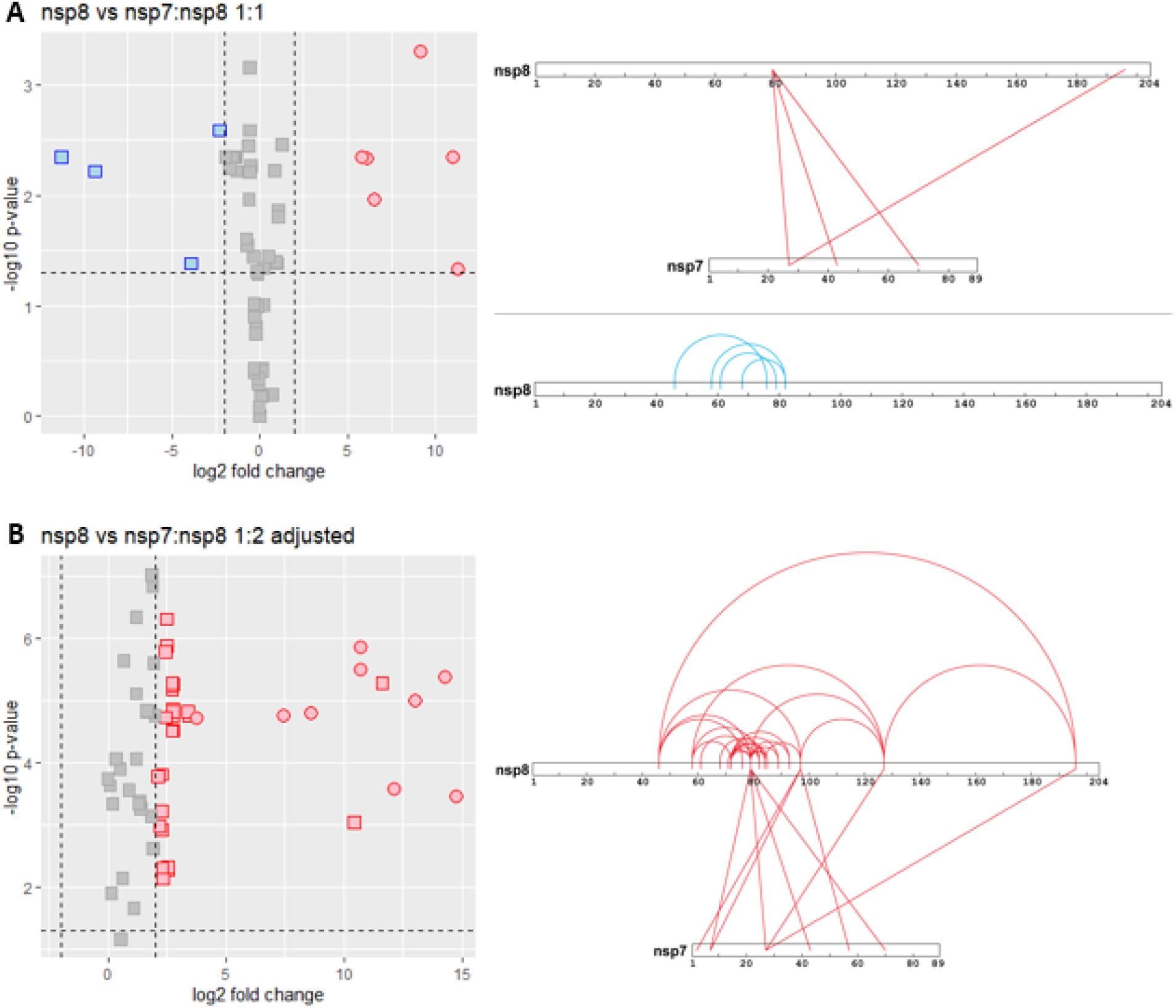
XL-MS relative quantification results of nsp7, nsp8, and nsp7:nsp8 complex. (A) Volcano plot summarizing the relative quantification results of crosslinks in nsp8 versus nsp7:nsp8 1:1. Crosslinks enriched in nsp7:nsp8 1:1 complex shown in red box and crosslinks enriched in the nsp8 alone sample shown in blue box. (B) Volcano plot summarizing the relative quantification results of nsp8 versus nsp7:nsp8 1:2 (adjusted) crosslinking reactions. Crosslinks enriched in nsp7:nsp8 1:1 complex shown in red box. Intra-nsp8 crosslinks shown as squares and inter-nsp7 nsp8 crosslinks shown as circles.

Previous studies have demonstrated that nsp7 and nsp8 play important roles as cofactors of nsp12 to give it processive RNA-dependent RNA polymerase activity [7]. Numerous cryo-EM structures of nsp7, nsp8, and nsp12 with or without RNA report a 1:1:2 nsp12:nsp7:nsp8 stoichiometry [6-10]. Based on these studies, we repeated the XL-MS analysis of the nsp7:nsp8 complex at a molar ratio of 1:2, respectively. For this differential analysis, the intensity values of intra-nsp8 crosslinks identified in the nsp7:nsp8 complex were halved in order to appropriately compare intensities to the nsp8 alone sample (**Figure 3B**). This analysis revealed four additional inter-nsp7-nsp8 crosslinks. While the majority of intra-nsp8 crosslinks remain unchanged between the two samples, some are slightly enriched in the nsp7:nsp8 complex when at a molar ratio of 1:2. The presence of these enriched intra-nsp8 crosslinks suggests that nsp8 interacts with nsp7 and nsp8 simultaneously, consistent with a 1:2 nsp7:nsp8 stoichiometry. Furthermore, nsp8 self-interaction has also been characterized by yeast two-hybrid screen and co-immunoprecipitation experiment [33] as well as novel phase separation-based method used to identify protein-protein interactions *in vitro* [34].

### HDX-MS and XL-MS results overlay to confirm nsp7:nsp8 interaction in solution

Overlaying the HDX-MS and XL-MS results onto the nsp7 and nsp8 sequences as shown in **Figure 4** demonstrates the complementary nature of the two techniques as the inter-nsp7-nsp8 crosslinks map to regions protected from solvent exchange in the differential HDX-MS experiment. Interestingly, regions containing both nsp8 Lys79 and Lys96, located within H1^nsp8^, demonstrate both the largest magnitude of protection from deuterium exchange and make the most inter-protein crosslinks with nsp7. These observations are consistent with the X-ray structure (PDB: 6YHU) showing H1^nsp8^ to be involved in stabilizing both the dimer and heterotetramer interface. An additional crosslink observed between nsp8 Lys196 (truncated in the aforementioned crystal structure) and nsp7 Lys27 suggests that additional contact sites are likely outside of the proposed helical bundle interactions.

**Figure 4.**
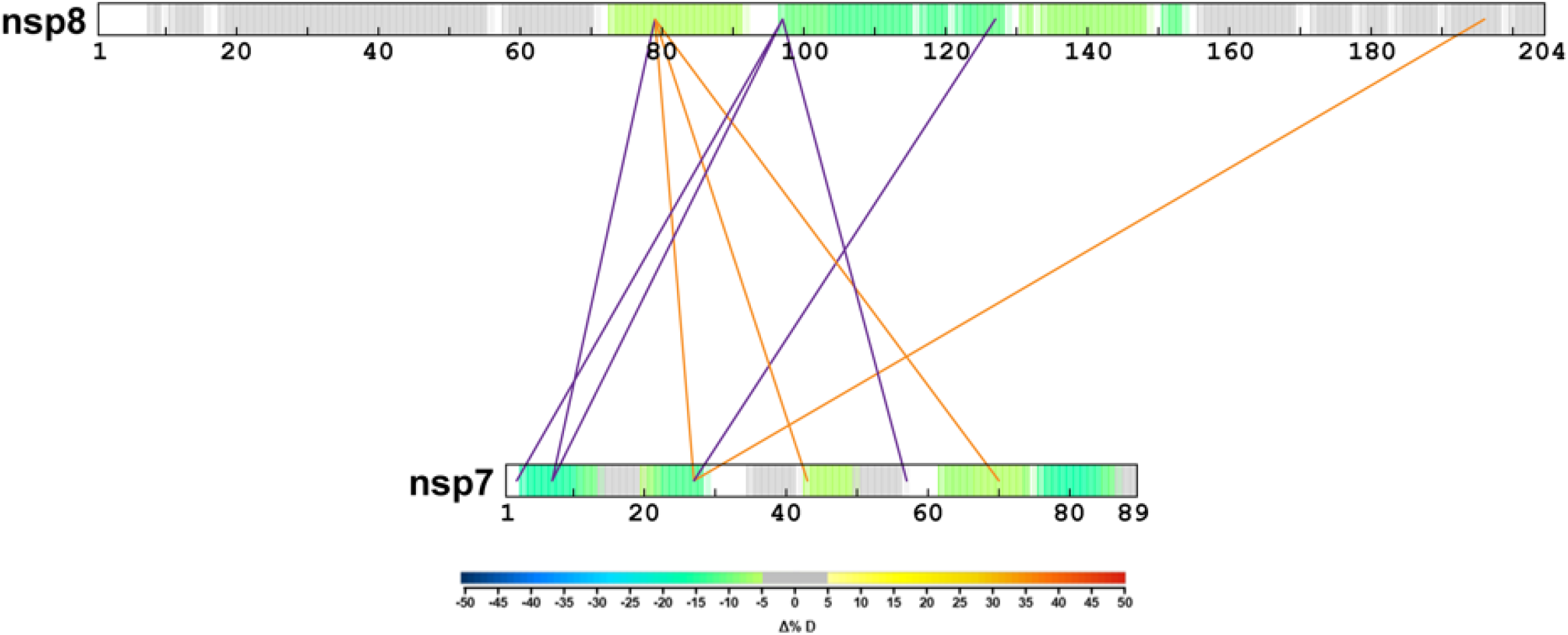
Overlay of HDX-MS and XL-MS results on nsp7 and nsp8 sequences. Observed inter-nsp7 nsp8 crosslinks only identified in the nsp7:nsp8 1:1 complex are colored in orange and inter-nsp7 nsp8 crosslinks also identified in nsp7:nsp8 1:2 complex are colored in blue. Consolidated change in percent deuterium uptake from Fig 1C and 1D are overlayed on the nsp7 and nsp8 sequences, respectively. Initial Met residue removed from both nsp7 and nsp8 sequences to maintain correct residue numbering.

### Mapping inter-nsp7-nsp8 crosslinks can be used to assess three dimensional structures

XL-MS analysis can be used to assess available three-dimensional structures and their relevance to solution-based structure. While the DSSO crosslinker measures 10.1 Å, it has a theoretical upper limit of approximately 26 Å to account for the orientation of side chains from crosslinked residues [35]. Xlink Analyzer [36] in Chimera [37] was used to map the inter-protein crosslinks onto all of the available nsp7:nsp8 crystal structures (**Table S3**). The calculated distances of the inter-protein crosslinks at either the dimer or heterotetramer interface within each group of structures are in good agreement. Comparing the inter-protein crosslinks mapped to the dimer interface between the three groups, good agreement in distances was observed. However, when comparing the inter-protein crosslinks mapped to the heterotetramer interface, differences are observed in which crosslinks are above or below the 26 Å limit. Furthermore, the cubic heterotetramer structure is unable to successfully map two crosslinks (nsp7 Lys7 to nsp8 Lys79 and nsp7 Lys43 to nsp8 Lys79) to either dimer or heterotetramer interfaces. On the contrary, all the inter-protein crosslinks were successfully mapped to either interface in the linear heterotetramer structures. This suggests that the linear heterotetramer structure is a better representation of the structural conformation of the nsp7:nsp8 complex in solution.

Focusing on the published nsp7:nsp8 structure (PDB: 6YHU) that shows the relevant linear heterotetramer conformation, we can map 10 of the 11 inter-protein crosslinks onto the structure (**Figure 5**). The crosslink identified to nsp8 Lys196 could not be mapped as this residue fell outside the truncated protein sequence used to obtain the structure. All 10 crosslinks were successfully mapped to either the dimer (**Figure 5A, Table 1: dimer interface**) or heterotetramer interface (**Figure 5B, Table 1: heterotetramer interface**) in PDB:6YHU. Only two crosslinks were successfully mapped to both interfaces: nsp7 Lys7 to nsp8 Lys97 and nsp7 Lys2 to nsp8 Lys97. These crosslinked residues are located within H1^nsp7^ and H1^nsp8^ which are the two helices predicted to be involved in forming both the dimer and heterotetramer interface. The third crosslink between these two helices (nsp7 Lys7 and nsp8 Lys79) has a calculated distance of 26.8 Å which is outside the threshold for DSSO distance of 26 Å. However, this threshold is not a hard cutoff as it tries to account for the various possible orientations of the Lys side chains. As such, the nsp7 Lys7 and nsp8 Lys79 crosslink should also be considered to be in agreement with the structure. Thus, all solution based inter-protein crosslinks observed between H1^nps7^ and H1^nsp8^ can be mapped to both the dimer and heterotetramer surfaces, and the remaining crosslinks can be mapped to either the dimer or heterotetramer interaction surfaces.

**Table 1.**
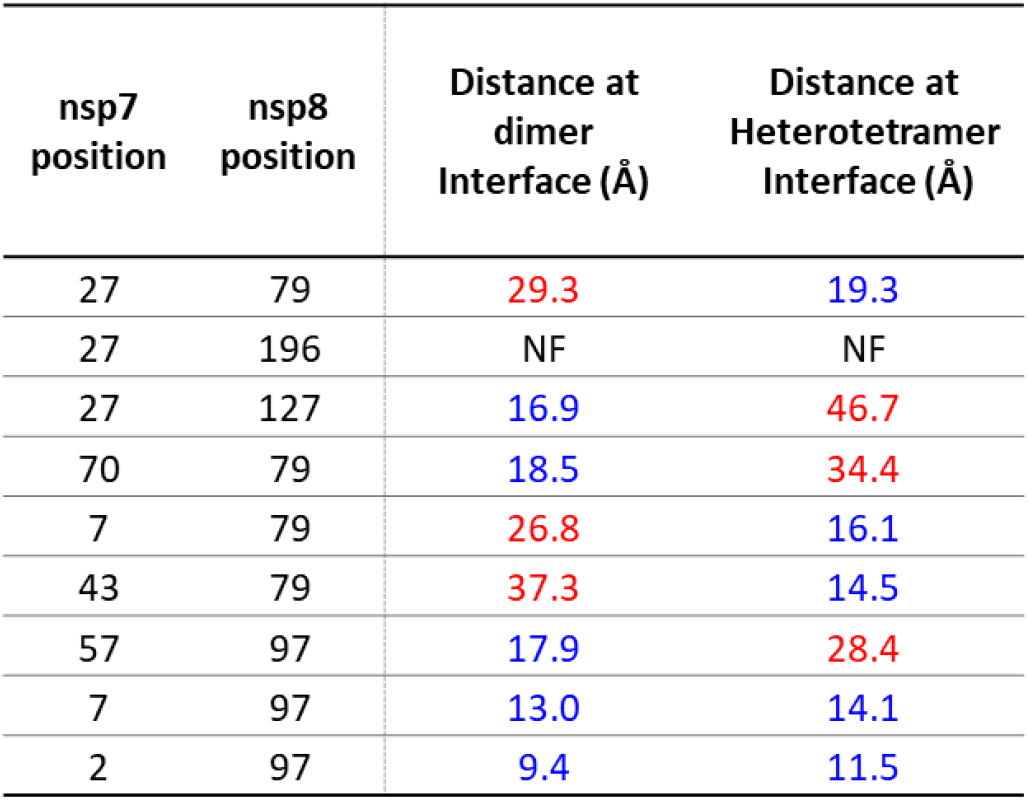
Distances of inter-nsp7 nsp8 crosslinks mapped to SARS-CoV-2 nsp7:nsp8 heterotetramer structure (PDB: 6YHU).

| nsp7 position | nsp8 position | Distance at dimer Interface (Å) | Distance at Heterotetramer Interface (Å) |
| --- | --- | --- | --- |
| 27 | 79 | 29.3 | 19.3 |
| 27 | 196 | NF | NF |
| 27 | 127 | 16.9 | 46.7 |
| 70 | 79 | 18.5 | 34.4 |
| 7 | 79 | 26.8 | 16.1 |
| 43 | 79 | 37.3 | 14.5 |
| 57 | 97 | 17.9 | 28.4 |
| 7 | 97 | 13.0 | 14.1 |
| 2 | 97 | 9.4 | 11.5 |

**Figure 5.**
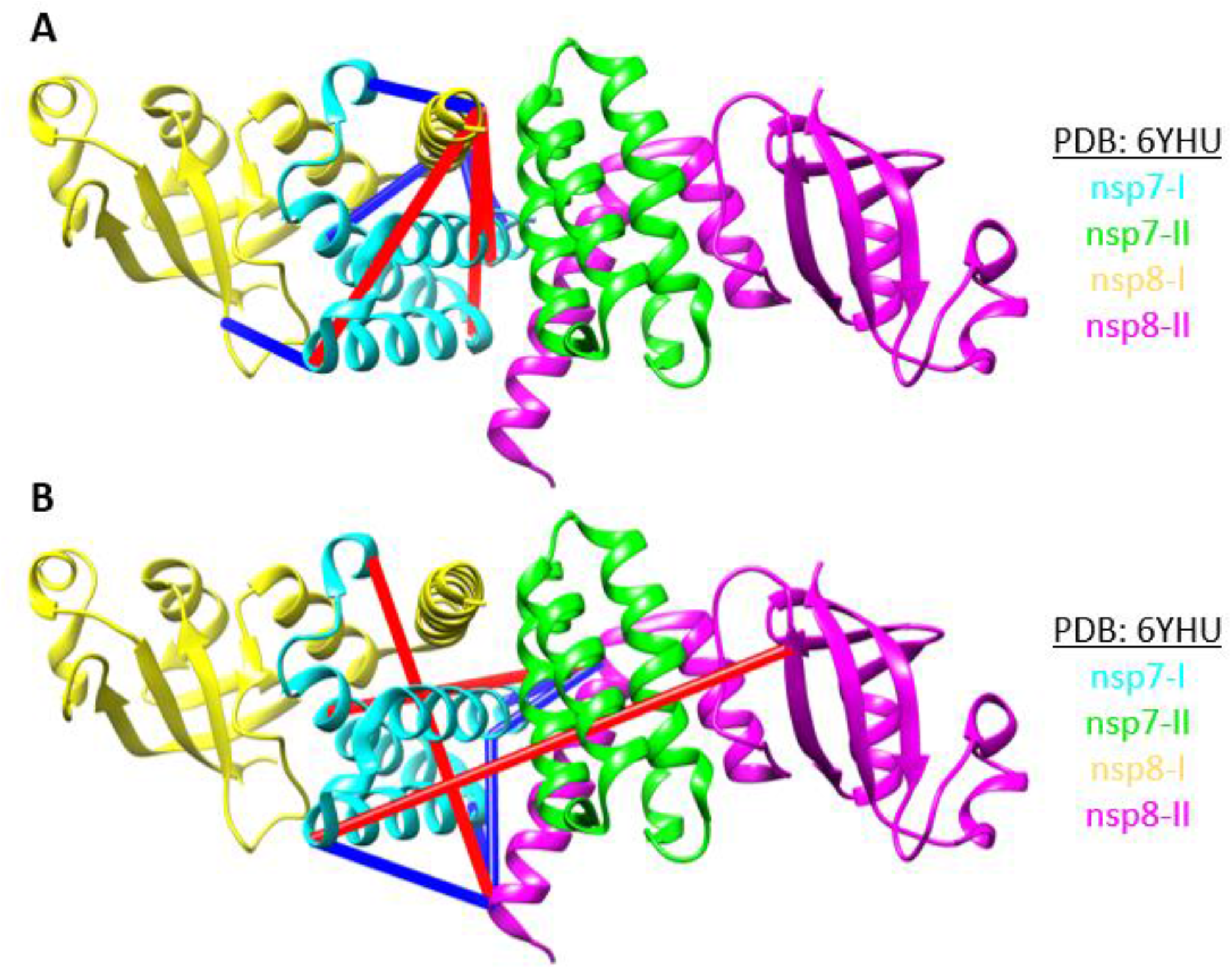
Mapping of nsp7 nsp8 inter-protein crosslinks on the dimer (A) and heterotetramer (B) interaction surfaces of the SARS-CoV-2 nsp7 nsp8 heterotetramer crystal structure (PDB: 6YHU). Crosslinks greater than 26 Å distance labeled in red and crosslinks less than 26 Å labeled in blue.

### Mapping inter-nsp7-nsp8 crosslinks to higher order assembly structures provides insight into RTC complex formation

The CoV-2 nsp7:nsp8 dimer was reported to superimpose well with the subunits of higher order assemblies of SARS-CoV nsp7:nsp8 and the cryo-EM structure of the replicating polymerase nsp7, nsp8, nsp12, and RNA [12]. To validate this finding, we mapped our inter-nsp7-nsp8 crosslinks onto these additional structures. Zhai *et al*. reported an hexadecameric circular complex for SARS-CoV nsp7:nsp8 [19]. They showed that nsp8 can exist in two different conformations, with one adopting a ‘golf-club’-like structure (**Figure 6: nsp8-II in pink**) and the other adopting a ‘golf-club’ with a bent shaft (**Figure 6: nsp8-I in yellow**). Nevertheless, the contacts between either nsp8 conformations and nsp7 are roughly the same. The nsp8 head is primarily responsible for interacting with nsp7 and this region is similar in either conformations. As shown in Figure 5A, mapping the observed inter-nsp7-nsp8 crosslinks to either possible SARS-CoV nsp7:nsp8 dimers shows similar crosslink distances in both conformations (**Table 2: dimer interface)**. These inter-protein crosslink distances from the SARS-CoV nsp7:nsp8 dimer interface are similar to the observed crosslink distances in the CoV-2 dimer (PDB:6YHU) (**Figure 6A, Table 2: dimer interface**). Of the 10 inter-nsp7-nsp8 crosslinks mapped in the SARS-CoV-2 structure, only one crosslink could not be mapped in the structure, as the corresponding residue of Lys70 is Arg70 in SARS-CoV. Comparing distances from the CoV-2 heterotetramer interface to the equivalent interface in SARS-CoV, we also observe similar crosslink distances (**Figure 6B, Table 2: heterotetramer interface**). However, when the crosslinks are mapped to the hexadecameric interface, all but one crosslink violate the 26 Å limit (**Figure 6C, Table 2: hexadecamer interface**). This suggests that full-length nsp7 and nsp8 in CoV-2 likely do not assemble into the higher order complex observed in CoV nsp7:nsp8.

**Table 2.**
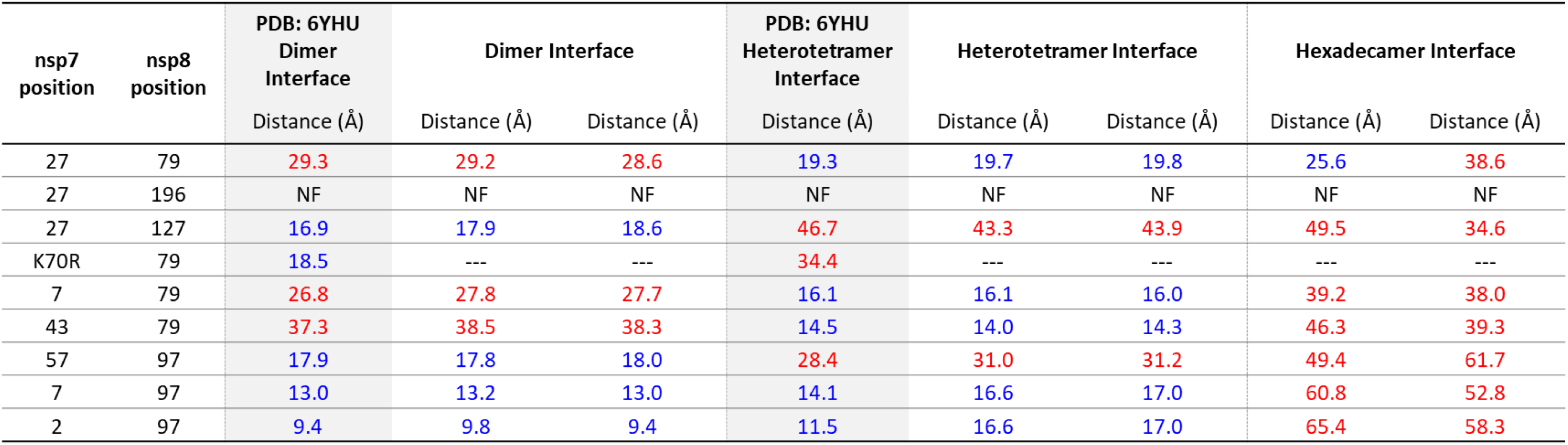
Distances of inter-nsp7 nsp8 crosslinks mapped to SARS-CoV nsp7:nsp8 hexadecamer structure (PDB: 2AHM).

**Figure 6.**
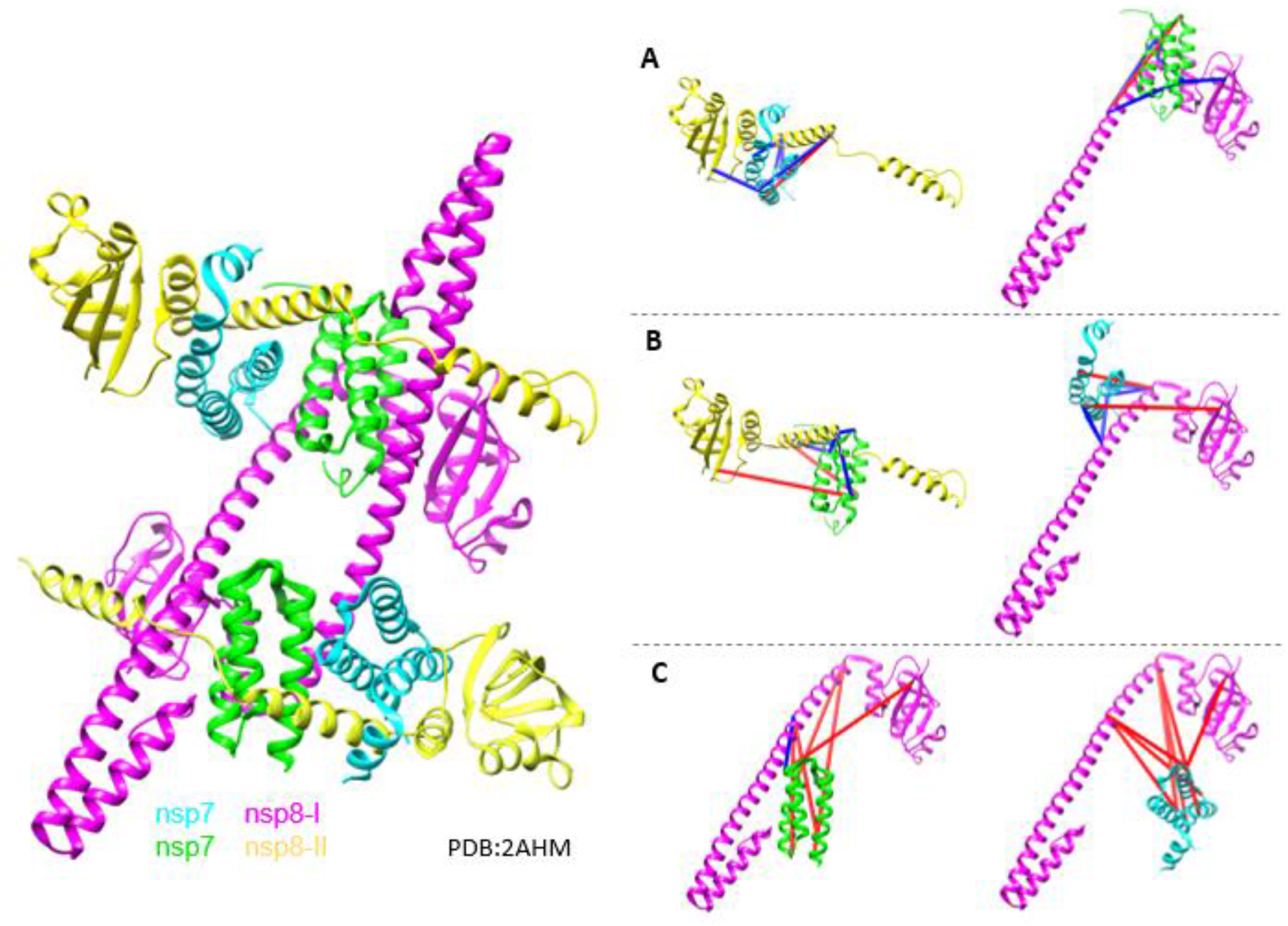
Mapping of nsp7 nsp8 inter-protein crosslinks on the dimer (A), heterotetramer (B), hexadecamer (C) interaction surfaces of the SARS-CoV nsp7 nsp8 hexadecamer cryo-EM structure (PDB: 2AHM). Crosslinks greater than 26 Å distance labeled in red and crosslinks less than 26 Å labeled in blue.

This finding is at odds with the proposed transition model by Wang *et al*. of the CoV-2 primase complex to the polymerase complex [8]. Their model suggests that RNA binds the hexadecameric nsp7-nsp8 structure to *de novo* synthesize primers before dissociating in halves to assemble with nsp12 pre-bound with a nsp8 molecule. The results presented here suggest that CoV-2 does not assemble into a hexadecameric structure, and thus would require RNA to be able to bind the nsp7:nsp8 dimer or heterotetramer to support primer *de novo* synthesis. Electrostatic surface analysis by Konkolova *et al*. suggests that while the dimer does not form any highly positively charged surfaces, the heterotetramer interface is framed by positively charged residues that can serve as the putative RNA-binding site [12]. We can thus hypothesize that RNA binds the heterotetramer structure to *de novo* synthesize primers and then dissociates into the dimers to allow assembly with nsp12. Additional experiments to confirm that the nsp7:nsp8 heterotetrameric structure can bind RNA as well as probe the structural dynamics of nsp7:nsp8:nsp12 and nsp7:nsp8:nsp12:RNA complexes in solution would be required to fully validate the mechanism of formation of the CoV-2 catalytic polymerase complex from the primase complex.

To investigate if the nsp7-nsp8 dimer can bind nsp12 in the given conformation, we mapped the inter-nsp7-nsp8 crosslinks to the replicating polymerase structure (PBD:6YYT) reported by Hillen *et al*. as the nsp7:nsp8 X-ray dimer was shown to superimpose well onto the nsp7:nsp8 dimer in this complex [7]. All 10 crosslinks from the nsp7:nsp8 structure were mapped and similar crosslink distances were determined (**Figure 7A, Table 3**). This suggests that nsp7:nsp8 binding to nsp12 does not alter nsp7:nsp8 interaction. Mapping the crosslinks between nsp7 and the additional nsp8 subunit gave crosslink distances all greater than 26 Å, consistent with the notion that the second nsp8 subunit does not make any contact with nsp7 and may be pre-bound to nsp12 (**Figure 7B, Table 3**).

**Table 3.** Distances of inter-nsp7 nsp8 crosslinks mapped to SARS-CoV-2 replicating polymerase structure (PDB: 6YYT).

| nsp7 position | nsp8 position | PDB: 6YHU<br>Dimer interface<br>Distance (Å) | PDB: 6YYT<br>nsp7-nsp8i<br>Distance (Å) | PDB: 6YYT<br>nsp7-nsp8ii<br>Distance (Å) |
| --- | --- | --- | --- | --- |
| 27 | 79 | 29.3 | 30.9 | 43.6 |
| 27 | 196 | NF | NF | NF |
| 27 | 127 | 16.9 | 17.6 | 27.7 |
| 70 | 79 | 18.5 | 22.1 | 52.5 |
| 7 | 79 | 26.8 | 23.4 | 48.5 |
| 43 | 79 | 37.3 | 33.9 | 55.5 |
| 57 | 97 | 17.9 | 17.7 | 48.9 |
| 7 | 97 | 13.0 | 13.3 | 51.0 |
| 2 | 97 | 9.4 | 10.3 | 59.4 |

**Figure 7.**
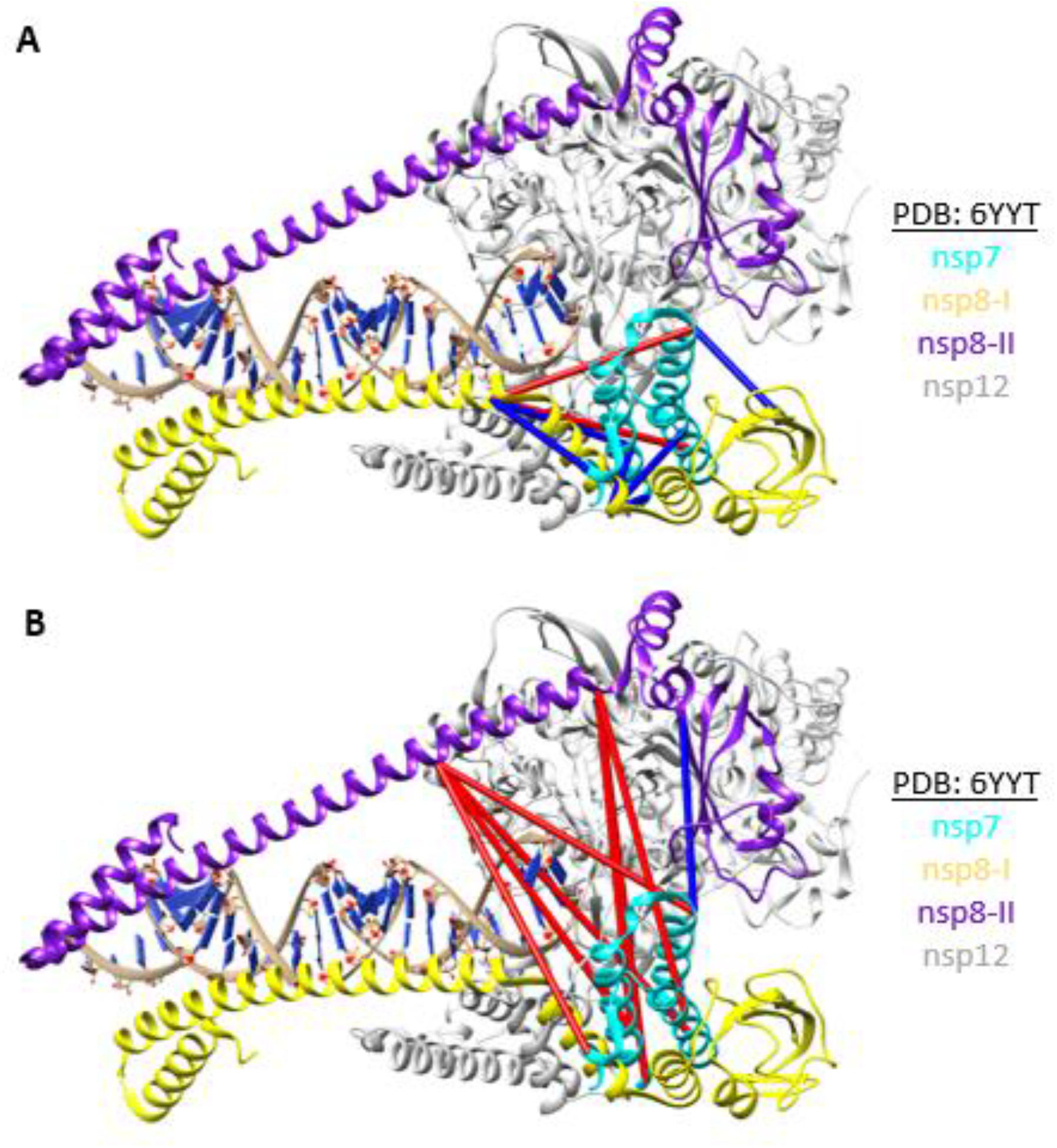
Mapping of nsp7-nsp8 inter-protein crosslinks to nsp8-I (A) or nsp8-II (B) of the SARS-CoV-2 replicating polymerase structure (PDB: 6YYT). Crosslinks greater than 26 Å distance labeled in red and crosslinks less than 26 Å labeled in blue.

Additional structures of the replicating polymerase have been reported since the structure published by Hillen *et al*., including structures with and without bound RNA. The other structure with RNA bound (PDB:7C2K) reports similar distances for the inter-nsp7-nsp8 crosslinks demonstrating reproducibility and generalizability between published structures (**Table 4**). The structures of the replicating polymerase without RNA bound (PDB: 7BV1, 7BW4, 7BTF, and 6M71) are unable to resolve as much of the nsp8 N-terminus and thus the crosslinks to nsp8 Lys80 cannot be mapped in these structures. All other inter-nsp7-nsp8 crosslinks were successfully mapped to the nsp7:nsp8 dimer and gave similar crosslink distances to the distances observed in the replicating polymerase with RNA bound (PDB:6YYT) or the dimer interface of the nsp7:nsp8 heterotetramer (PDB:6YHU) (**Table 5**). These observations suggest that presence of RNA does not alter nsp7:nsp8 interaction in complex with nsp12, in concordance to what can be observed by the superposition of apo and RNA-bound cryo-EM structures. Additional XL-MS and HDX-MS experiments with nsp7, nsp8, nsp12, and RNA would be required to further validate the transition model of CoV-2 primase complex to replicating polymerase complex that facilitates viral RNA synthesis.

**Table 4.**
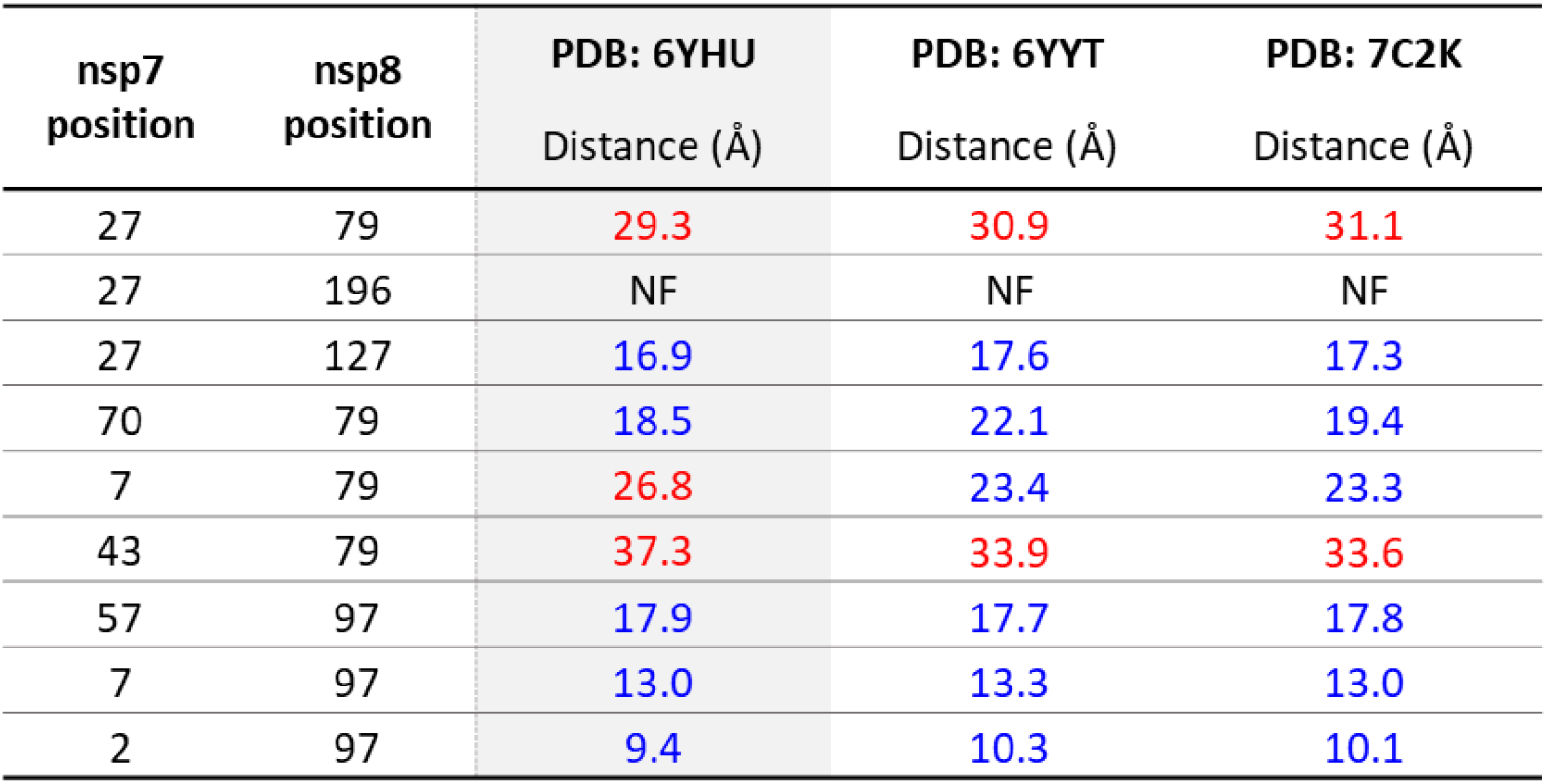
Distances of inter-nsp7 nsp8 crosslinks mapped to available SARS-CoV-2 replicating polymerase structures composed of nsp7, nsp8, nsp12, and RNA.

| nsp7 position | nsp8 position | PDB: 6YHU<br>Distance (Å) | PDB: 6YYT<br>Distance (Å) | PDB: 7C2K<br>Distance (Å) |
| --- | --- | --- | --- | --- |
| 27 | 79 | 29.3 | 30.9 | 31.1 |
| 27 | 196 | NF | NF | NF |
| 27 | 127 | 16.9 | 17.6 | 17.3 |
| 70 | 79 | 18.5 | 22.1 | 19.4 |
| 7 | 79 | 26.8 | 23.4 | 23.3 |
| 43 | 79 | 37.3 | 33.9 | 33.6 |
| 57 | 97 | 17.9 | 17.7 | 17.8 |
| 7 | 97 | 13.0 | 13.3 | 13.0 |
| 2 | 97 | 9.4 | 10.3 | 10.1 |

**Table 5.**
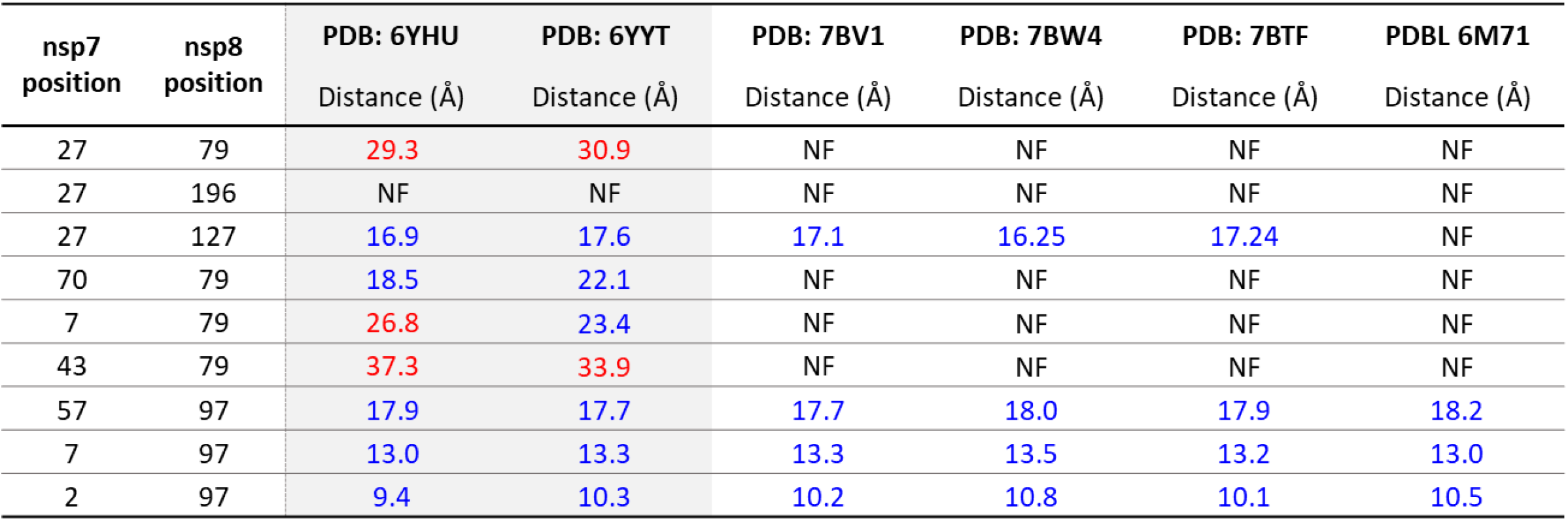
Distances of inter-nsp7 nsp8 crosslinks mapped to available SARS-CoV-2 replicating polymerase structures composed of nsp7, nsp8, and nsp12.

| nsp7<br>position | nsp8<br>position | PDB: 6YHU<br>Distance (Å) | PDB: 6YYT<br>Distance (Å) | PDB: 7BV1<br>Distance (Å) | PDB: 7BW4<br>Distance (Å) | PDB: 7BTF<br>Distance (Å) | PDBL 6M71<br>Distance (Å) |
| --- | --- | --- | --- | --- | --- | --- | --- |
| 27 | 79 | 29.3 | 30.9 | NF | NF | NF | NF |
| 27 | 196 | NF | NF | NF | NF | NF | NF |
| 27 | 127 | 16.9 | 17.6 | 17.1 | 16.25 | 17.24 | NF |
| 70 | 79 | 18.5 | 22.1 | NF | NF | NF | NF |
| 7 | 79 | 26.8 | 23.4 | NF | NF | NF | NF |
| 43 | 79 | 37.3 | 33.9 | NF | NF | NF | NF |
| 57 | 97 | 17.9 | 17.7 | 17.7 | 18.0 | 17.9 | 18.2 |
| 7 | 97 | 13.0 | 13.3 | 13.3 | 13.5 | 13.2 | 13.0 |
| 2 | 97 | 9.4 | 10.3 | 10.2 | 10.8 | 10.1 | 10.5 |

## CONCLUSIONS

HDX-MS and XL-MS are complementary structural proteomic techniques used to characterize proteins and protein complexes. While X-ray crystallography and cryo-EM present proteins in one or more predominant conformations, HDX-MS and XL-MS interrogate proteins in-solution to allow for direct analysis of protein conformational dynamics. HDX-MS specifically probes protein backbone dynamics while XL-MS reports on side-chain residency and reactivity. This information can then be used to interrogate models of three-dimensional structures and ultimately inform how protein structure relates to biological function.

Herein we used HDX-MS and XL-MS to probe the solution phase structural dynamics of the CoV-2 nsp7:nsp8 complex and assess the relevance of the available crystal structures. While the crystal structures suggest three possible structural conformations (dimer, linear heterotetramer, and cubic heterotetramer), the HDX-MS and XL-MS results obtained suggest that the linear heterotetramer structure is likely to be the most relevant structure for the complex in solution. This structure, represented by the published structure PDB: 6YHU, proposes a 2:2 heterotetrameric structure formed via interaction of H1^nsp7^ and H3^nsp7^ with H1^nsp8^ and H2^nsp8^ at the dimer interface and H1^nsp7^ and H2^nsp7^ with H1^nsp8^ at the heterotetramer interface. HDX-MS revealed protection from exchange in these regions and XL-MS identified inter-nsp7-nsp8 crosslinks that were mapped to these protected regions. Furthermore, all inter-protein crosslinks were below the 26 Å distance limit when mapped to either the dimer or heterotetramer interface of this nsp7:nsp8 2:2 heterotetrameric structure. Additional protection to exchange at the C-terminus of nsp7, as well as a crosslink from H1^nsp7^ to the C-terminus of nsp8, suggests that additional contact surfaces are likely to be present, involving the N- and C-termini of nsp7 and nsp8, that are not present in the PDB structures due to the use of truncated proteins for crystallization or lack of electron density.

## Acknowledgements

Molecular graphics and analyses performed with UCSF Chimera, developed by the Resource for Biocomputing, Visualization, and Informatics at the University of California, San Francisco, with support from NIH U54 AI150472 to E.A. and P.R.G. The E.A. laboratory is also grateful for financial support from a Rutgers Center for COVID-19 Response and Pandemic Preparedness COVID research award.

