## Supplemental figures and tables for "Resolving the Dynamic Motions of SARS-CoV-2 nsp7 and nsp8 Proteins Using Structural Proteomics"

**Table S1.** nsp8 peptides sequences shown with their corresponding charge state and residue start and end number based on sequence. The perturbation values Δ%D reported with propagated errors are shown in parenthesis.

| nsp8 Sequence | Charge | Start | End | Features | nsp8 vs nsp7nsp8 1:1 | nsp8 vs nsp7nsp8 3:1 |
|---|---|---|---|---|---|---|
| FSSLPSY | 1 | 6 | 12 | | -1 (3) | 1 (2) |
| FSSLPSYAAF | 2 | 6 | 15 | | -1 (3) | 0 (2) |
| ATAQEAYEQAVANGDS | 2 | 16 | 31 | | 4 (2) | 0 (2) |
| ATAQEAYEQAVANGDSE | 2 | 16 | 32 | | 5 (2) | 5 (2) |
| ATAQEAYEQAVANGDSEVVLKKLKKSLNVAKSEFDRDAAM | 4 | 16 | 55 | | 3 (2) | 5 (1) |
| YEQAVANGDSE | 2 | 22 | 32 | | -1 (2) | 0 (2) |
| YEQAVANGDSEVVLKKLKKSLNVAKSEFDRDAAM | 4 | 22 | 55 | | 3 (2) | 2 (2) |
| VVLKKLKKSLNVAKSEF | 2 | 33 | 49 | | 5 (3) | 3 (1) |
| VVLKKLKKSLNVAKSEF | 3 | 33 | 49 | | 5 (2) | 4 (3) |
| QRKLEKMAD | 2 | 56 | 64 | | 1 (3) | 4 (3) |
| QRKLEKMAD | 3 | 56 | 64 | | -1 (3) | 4 (3) |
| QRKLEKMADQAM | 3 | 56 | 67 | | 0 (3) | 1 (3) |
| QRKLEKMADQAMTQM | 2 | 56 | 70 | | -2 (3) | -2 (3) |
| QRKLEKMADQAMTQM | 3 | 56 | 70 | | 0 (3) | 1 (3) |
| QRKLEKMADQAMTQM | 4 | 56 | 70 | | 0 (3) | 0 (3) |
| YKQARSEDKRAKVTSAMQTML | 2 | 71 | 91 | [H1:14] | -2 (2) | -7 (2) |
| YKQARSEDKRAKVTSAMQTML | 3 | 71 | 91 | [H1:14] | -1 (3) | -6 (2) |
| YKQARSEDKRAKVTSAMQTML | 4 | 71 | 91 | [H1:14] | -1 (3) | -6 (2) |
| LRKLDNDAL | 3 | 95 | 103 | [H1:5 H2:2] | -3 (2) | -11 (3) |
| LRKLDNDALNNIINNARDGCVPLNIIPLTT | 4 | 95 | 124 | [H1:5 H2:12] | -7 (4) | -13 (4) |
| RKLDNDALNNIINNARDGCVPLNIIPLTT | 3 | 96 | 124 | [H1:4 H2:12] | -6 (2) | -15 (3) |
| RKLDNDALNNIINNARDGCVPLNIIPLTT | 4 | 96 | 124 | [H1:4 H2:12] | -6 (3) | -14 (3) |
| RKLDNDALNNIINNARDGCVPLNIIPLTTA | 4 | 96 | 125 | [H1:4 H2:12] | -6 (2) | -9 (2) |
| RKLDNDALNNIINNARDGCVPLNIIPLTTAAKL | 4 | 96 | 128 | [H1:4 H2:12 B1:3] | -6 (3) | -10 (2) |
| NNIINNARDGCVPLNIIPLTT | 2 | 104 | 124 | [H2:10] | -7 (3) | -13 (3) |
| NNIINNARDGCVPLNIIPLTTA | 2 | 104 | 125 | [H2:10] | -9 (3) | -14 (3) |
| NNIINNARDGCVPLNIIPLTTAAKL | 3 | 104 | 128 | [H2:10 B1:3] | -6 (2) | -8 (2) |
| MVVIPDYNT | 1 | 129 | 137 | [B1:5 H3:2] | 2 (3) | -7 (2) |
| MVVIPDYNTYKNTCDGTTF | 2 | 129 | 147 | [B1:5 H3:7 B2:1] | 3 (3) | -7 (2) |
| MVVIPDYNTYKNTCDGTTFT | 2 | 129 | 148 | [B1:5 H3:7 B2:2] | 2 (2) | -7 (2) |
| YASAL | 1 | 149 | 153 | [B2:2 B3:1] | -9 (4) | -10 (3) |
| WEIQQVVDADSKIVQL | 2 | 154 | 169 | [B3:8] | 4 (2) | -2 (3) |
| DSKIVQL | 1 | 163 | 169 | | 2 (2) | -2 (3) |
| DSKIVQL | 2 | 163 | 169 | | 2 (2) | -2 (3) |
| SEISM | 1 | 170 | 174 | | 0 (3) | 2 (3) |
| SEISMDNSPNLAWPLIVTAL | 2 | 170 | 189 | [B4:5] | 2 (1) | -1 (1) |
| ISMDNSPNLAWPLIVTAL | 2 | 172 | 189 | [B4:5] | 2 (1) | -2 (1) |
| DNSPNLAWPLIVT | 2 | 175 | 187 | [B4:3] | 5 (1) | -3 (1) |
| DNSPNLAWPLIVTA | 2 | 175 | 188 | [B4:4] | 4 (1) | 0 (2) |
| DNSPNLAWPLIVTAL | 2 | 175 | 189 | [B4:5] | 1 (1) | 1 (1) |
| LRANSA | 2 | 189 | 194 | [B4:3] | -2 (4) | 2 (3) |
| RANSAVKLQ | 2 | 190 | 198 | [B4:2] | 2 (4) | 1 (3) |
| QENLYFQ | 2 | 198 | 204 | [Tag:6] | -3 (2) | 4 (2) |

**Table S2.** nsp7 peptides sequences shown with their corresponding charge state and residue start and end number based on sequence. The perturbation values ∆%D reported with propagated errors are shown in parenthesis.

| nsp7 Sequence | Charge | Start | End | Features | nsp7 vs nsp7nsp8 1:1 | nsp7 vs nsp7nsp8 3:1 |
|---|---|---|---|---|---|---|
| SKMSDVKC | 2 | 1 | 8 | [H1:6] | -2 (3) | -20 (3) |
| SKMSDVKCTS | 1 | 1 | 10 | [H1:8] | -2 (3) | -15 (3) |
| SKMSDVKCTS | 2 | 1 | 10 | [H1:8] | -2 (4) | -15 (3) |
| SKMSDVKCTS | 3 | 1 | 10 | [H1:8] | -2 (4) | -16 (3) |
| SKMSDVKCTSV | 2 | 1 | 11 | [H1:9] | -3 (4) | -16 (3) |
| SKMSDVKCTSVVL | 2 | 1 | 13 | [H1:11] | -4 (4) | -14 (5) |
| SKMSDVKCTSVVL | 3 | 1 | 13 | [H1:11] | -4 (4) | -17 (4) |
| VLLSVL | 1 | 12 | 17 | [H1:6] | -3 (3) | 2 (4) |
| LSVLQQ | 1 | 14 | 19 | [H1:6] | 2 (2) | 3 (4) |
| LSVLQQL | 2 | 14 | 20 | [H1:7] | 0 (2) | 1 (2) |
| QQLRVESSSKL | 2 | 18 | 28 | [H1:4 H2:2] | 0 (3) | -7 (3) |
| QQLRVESSSKL | 3 | 18 | 28 | [H1:4 H2:2] | 0 (4) | -8 (3) |
| LRVESSSKL | 2 | 20 | 28 | [H1:2 H2:2] | -2 (4) | -14 (2) |
| RVESSSKL | 2 | 21 | 28 | [H1:1 H2:2] | -2 (4) | -17 (2) |
| VQLHNDIL | 1 | 33 | 40 | [H2:8] | 2 (2) | 4 (2) |
| VQLHNDIL | 2 | 33 | 40 | [H2:8] | 1 (2) | 4 (2) |
| VQLHNDILL | 2 | 33 | 41 | [H2:9] | 3 (2) | 4 (2) |
| HNDILL | 1 | 36 | 41 | [H2:6] | 3 (3) | 5 (3) |
| LAKDTTEA | 2 | 41 | 48 | [H2:2 H3:3] | -1 (4) | -8 (2) |
| LAKDTTEAF | 2 | 41 | 49 | [H2:2 H3:4] | -1 (3) | -8 (3) |
| AKDTTEAF | 2 | 42 | 49 | [H2:1 H3:4] | -1 (4) | -7 (4) |
| FEKMVSL | 1 | 49 | 55 | [H3:7] | 0 (3) | 5 (3) |
| FEKMVSL | 2 | 49 | 55 | [H3:7] | 1 (3) | 4 (3) |
| FEKMVSLL | 1 | 49 | 56 | [H3:8] | 0 (2) | 3 (2) |
| FEKMVSLL | 2 | 49 | 56 | [H3:8] | 0 (2) | 3 (2) |
| LSMQGAVDINKL | 1 | 60 | 71 | [H3:3] | 0 (3) | -7 (2) |
| LSMQGAVDINKLCE | 2 | 60 | 73 | [H3:3] | -2 (2) | -8 (3) |
| LSMQGAVDINKLCEE | 2 | 60 | 74 | [H3:3] | -2 (3) | -9 (3) |
| AVDINKL | 2 | 65 | 71 | | -2 (3) | -5 (3) |
| AVDINKLCE | 1 | 65 | 73 | | -2 (2) | -4 (3) |
| AVDINKLCE | 2 | 65 | 73 | [H4:7] | -2 (2) | -4 (3) |
| EMLDNRATL | 2 | 74 | 82 | [H4:6] | -2 (3) | -13 (3) |
| EMLDNRATLQENL | 2 | 74 | 86 | [H4:6 Tag:2] | -1 (4) | -11 (3) |
| MLDNRATL | 2 | 75 | 82 | [H4:5] | -2 (4) | -18 (2) |
| MLDNRATLQE | 2 | 75 | 84 | [H4:5] | -3 (4) | -17 (2) |
| MLDNRATLQENL | 2 | 75 | 86 | [H4:5 Tag:2] | -4 (5) | -14 (3) |
| LDNRATL | 2 | 76 | 82 | [H4:4] | -3 (4) | -18 (3) |
| LDNRATLQENL | 2 | 76 | 86 | [H4:4 Tag:2] | -1 (4) | -15 (3) |
| QENLYFQ | 1 | 83 | 89 | [Tag:5] | -3 (3) | -3 (2) |

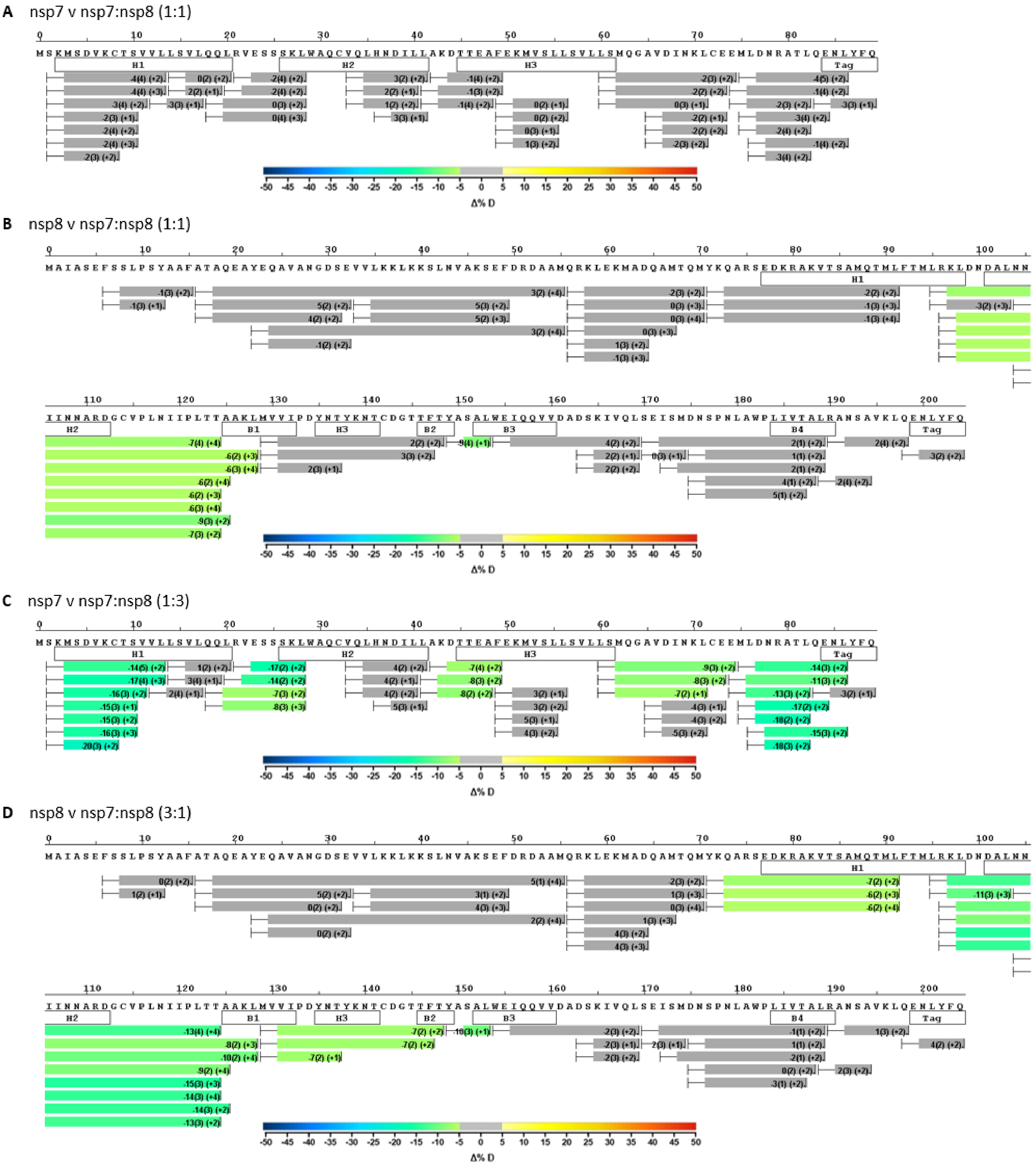


**Figure S1.** Sequence coverage for differential HDX-MS analysis of nsp7 vs nsp7:nsp8 1:1 (**A**), nsp8 vs nsp7:nsp8 1:1 (**B**), nsp7 vs nsp7:nsp8 1:3 (**C**), and nsp8 vs nsp7:nsp8 3:1 (**D**).

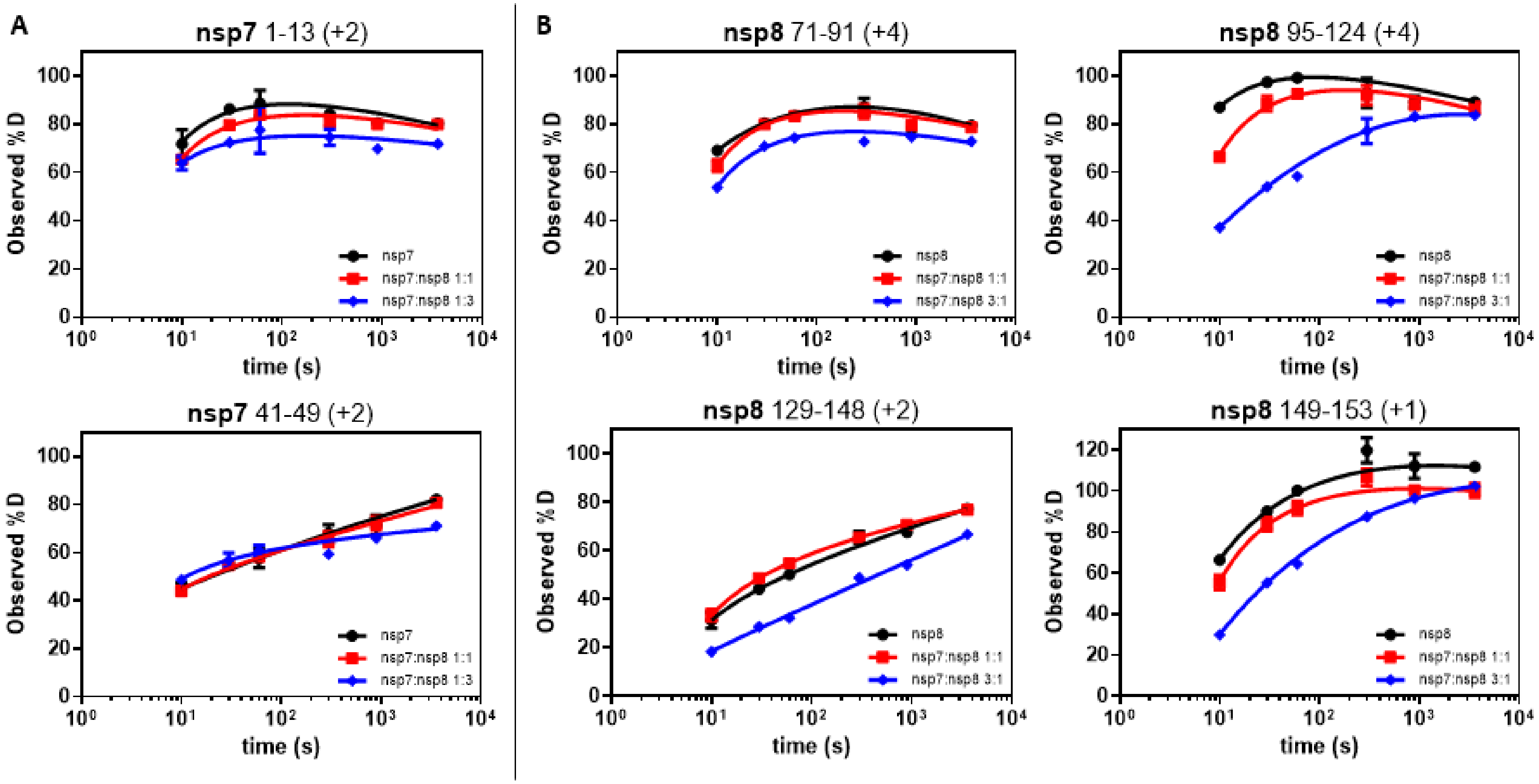


**Figure S2.** Representative deuterium build-up plots showing regions of protection in nsp7 (**A**) and nsp8 (**B**) upon complex formation.

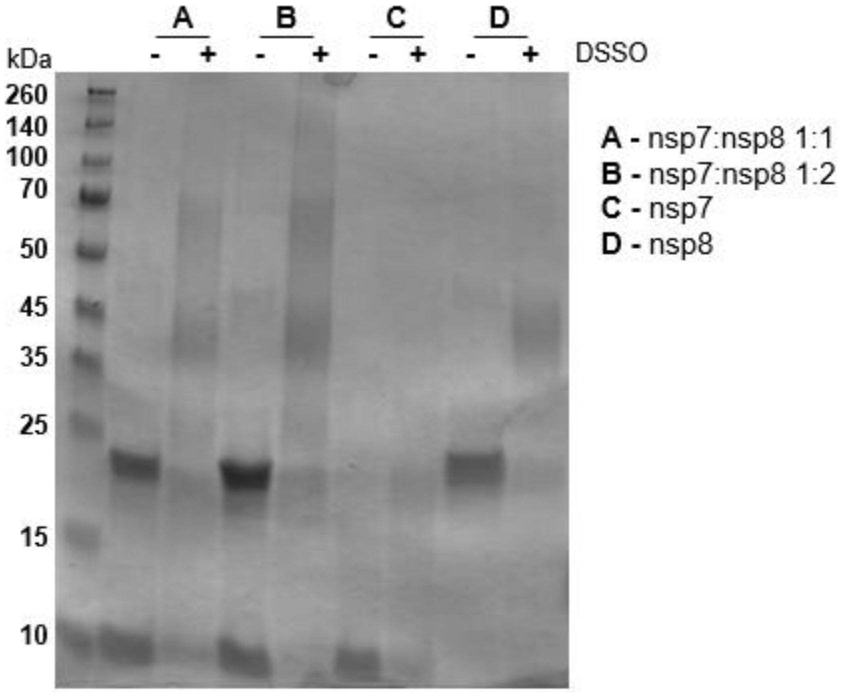


**Figure S3.** Crosslinking efficiency confirmed by SDS-PAGE. Presence of higher molecular weight bands and smeared bands in the sample with DSSO indicate successful crosslinking.

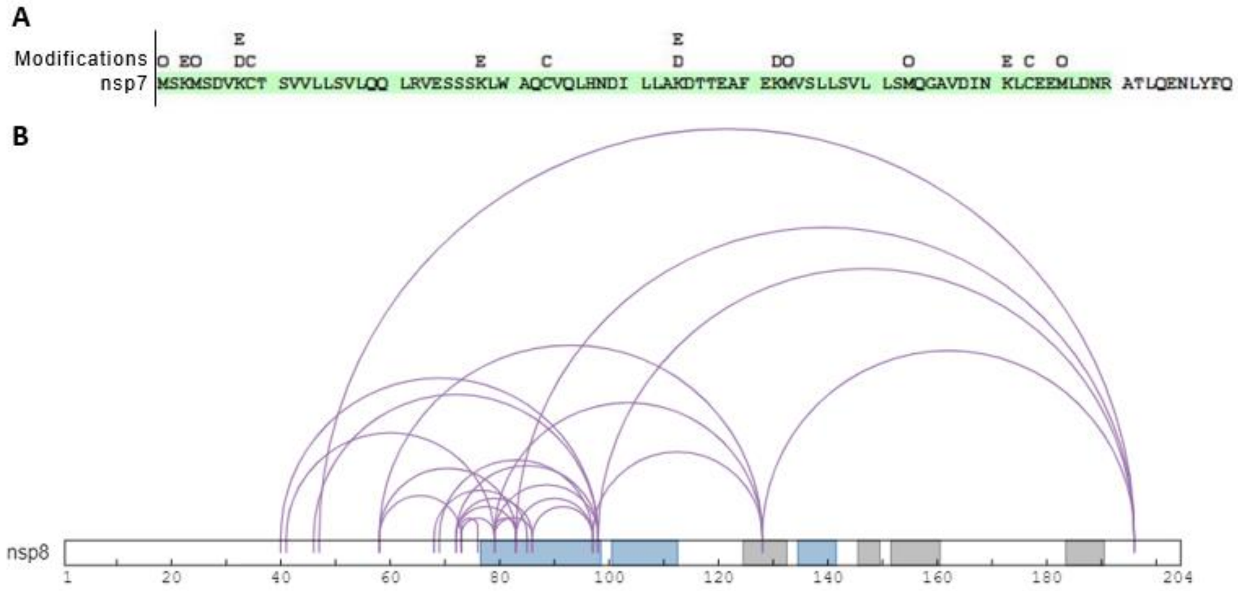


**Figure S4.** XL-MS of nsp7 and nsp8 XL in isolation. **(A)** Sequence coverage of nsp7 XL-MS with modifications annotated – oxidation (O), carbamidomethyl (C), DSSO Tris (D), DSSO hydrolyzed (E), and N-terminal acetylation (A). **(B)** nsp8 crosslinks mapped onto nsp8 sequence with secondary structure from PDB:6YHU annotated with α-helices in blue and β-sheets in gray. Initial Met residue removed from nsp8 sequence to maintain correct residue numbering.

**Table S2.** Distances of inter-nsp7 nsp8 crosslinks mapped to all available SARS-CoV-2 nsp7:nsp8 structures. Columns color coded by grouping of structures based on conformation: dimer (green), cubic heterotetramer (yellow), linear heterotetramer (gray). Measure crosslink distances reported in Å and labeled in red if they violate the 26 Å limit or in blue if they are below the 26 Å limit.

| nsp7 | nsp8 | Dimer Interface | | | | | | | | | | | | Heterotetramer Interface | | | | | | | | | |
|---|---|---|---|---|---|---|---|---|---|---|---|---|---|---|---|---|---|---|---|---|---|---|---|
| | | 6WIQ | 6M5I | 6WTC | | 6XIP | | 6WQD | | 7JLT | | 6YHU | | 6WTC | | 6XIP | | 6WQD | | 7JLT | | 6YHU | |
| | | (Å) | (Å) | (Å) | (Å) | (Å) | (Å) | (Å) | (Å) | (Å) | (Å) | (Å) | (Å) | (Å) | (Å) | (Å) | (Å) | (Å) | (Å) | (Å) | (Å) | (Å) | (Å) |
| 27 | 79 | 29.6 | 28.7 | 29.4 | 30.2 | 29.2 | 30.1 | 30.5 | 30.0 | 28.6 | 28.9 | 29.3 | 29.8 | 25.4 | 36.7 | 25.3 | 36.8 | 19.4 | 18.4 | 19.1 | 19.5 | 19.3 | 19.4 |
| 27 | 196 | NF | NF | NF | NF | NF | NF | NF | 33.7 | NF | NF | NF | NF | NF | NF | NF | NF | NF | 61.1 | NF | NF | NF | NF |
| 27 | 127 | 17.6 | 17.6 | 17.9 | 16.2 | 18.2 | 16.6 | 18.7 | 18.3 | 17.5 | 17.6 | 16.9 | 17.6 | 39.7 | 52.1 | 39.7 | 52.5 | 47.7 | 47.5 | 45.9 | 45.9 | 46.7 | 46.1 |
| 70 | 79 | 18.9 | 18.7 | 19.2 | 18.9 | 18.5 | 18.5 | 18.9 | 19.0 | 18.4 | 18.8 | 18.5 | 18.1 | 43.7 | 25.7 | 43.0 | 25.7 | 34.6 | 33.6 | 37.5 | 38.2 | 34.4 | 33.9 |
| 7 | 79 | 27.8 | 27.7 | 27.3 | 27.1 | 27.1 | 26.8 | 27.6 | 26.8 | 27.6 | 27.5 | 26.8 | 26.7 | 47.2 | 31.1 | 47.0 | 31.2 | 17.2 | 16.3 | 15.8 | 16.1 | 16.1 | 16.4 |
| 43 | 79 | 38.7 | 38.5 | 38.4 | 37.7 | 38.3 | 37.3 | 38.3 | 37.9 | 38.2 | 38.4 | 37.3 | 37.5 | 50.5 | 32.6 | 50.1 | 32.7 | 15.6 | 15.3 | 14.2 | 14.7 | 14.5 | 14.9 |
| 57 | 97 | 18.1 | 18.0 | 18.0 | 17.9 | 18.0 | 17.9 | 17.9 | 17.9 | 17.9 | 17.8 | 17.9 | 17.9 | 21.5 | 43.9 | 21.8 | 44.1 | 28.3 | 28.6 | 28.6 | 28.2 | 28.4 | 28.6 |
| 7 | 97 | 13.4 | 13.2 | 13.0 | 13.1 | 13.1 | 13.0 | 12.8 | 13.0 | 13.2 | 13.5 | 13.0 | 12.8 | 34.3 | 45.8 | 34.5 | 45.9 | 13.7 | 14.0 | 14.9 | 14.7 | 14.1 | 14.0 |
| 2 | 97 | 10.5 | 10.0 | 9.5 | 9.0 | 9.4 | 9.5 | 9.4 | 9.1 | 9.9 | 10.0 | 9.4 | 9.1 | 39.0 | 39.1 | 39.0 | 39.4 | 11.4 | 11.3 | 12.7 | 13.0 | 11.5 | 11.3 |
